# Rational engineering of *Kluyveromyces marxianus* to create a chassis for the production of aromatic products

**DOI:** 10.1101/2020.08.12.247957

**Authors:** Arun S. Rajkumar, John P. Morrissey

## Abstract

The yeast *Kluyveromyces marxianus* offers unique potential for industrial biotechnology because of useful features like rapid growth, thermotolerance and a wide substrate range. As an emerging alternative platform, *K. marxianus* requires the development and validation of metabolic engineering strategies to best utilize its metabolism as a basis for bio-based production. To illustrate the synthetic biology strategies to be followed and showcase its potential, we describe a comprehensive approach to rationally engineer a metabolic pathway in *K. marxianus*. We use the phenylalanine biosynthetic pathway both as a prototype and because phenylalanine is a precursor for commercially valuable secondary metabolites. First, we modify and overexpress the pathway to be resistant to feedback inhibition so as to overproduce phenylalanine *de novo* from synthetic minimal medium. Second, we assess native and heterologous means to increase precursor supply to the biosynthetic pathway. Finally, we eliminate branch points and competing reactions in the pathway and rebalance precursors to redirect metabolic flux to a specific product, 2-phenylethanol (2-PE). As a result, we are able to construct robust strains capable of producing over 800 mg L^−1^ 2-PE from minimal medium. The strains we constructed are a promising platform for the production of aromatic amino acid-based biochemicals, and our results illustrate challenges with attempting to combine individually beneficial modifications in an integrated platform.

## 1. Introduction

Microbial cell factories are an important part of an emerging sustainable bioeconomy. By metabolically engineering microbial hosts, it is possible to synthesize chemical compounds that are currently sourced from non-renewable resources or from renewable resources that may not meet the growing demands of the chemical, food and pharmaceutical industries. The process typically involves cloning heterologous pathways for such compounds into host microbes and altering native pathways to ensure that metabolism is optimized for the products of interest. The budding yeast *Saccharomyces cerevisiae* has proved to be an excellent host for *de novo* synthesis of valuable secondary metabolites in the flavonoid, stilbenoid and alkaloid families (Liu et al., 2020). Several of these are directly derived from aromatic amino acids: flavonoids/stilbenoids and aroma compounds from phenylalanine (Koopman et al., 2012); benzylisquinoline alkaloids from tyrosine (Li et al., 2018); and monoterpene alkaloids from tryptophan (Brown et al., 2015). Others are derived from intermediates of the shikimate pathway, such as the flavour compound vanillin (Strucko et al., 2015) and the fine chemical muconic acid (Curran et al., 2013). Optimizing precursor supply is of special interest if these compounds are to be produced at titres for commercial production competitive with existing commercial sources. As a result, the metabolic engineering of *S. cerevisiae* to overproduce aromatic amino acids (AAAs) has been the subject of considerable research and development to produce diverse derived secondary metabolites of commercial value (Gottardi et al., 2017; Liu et al., 2020). Furthermore, phenylalanine and tyrosine are commercial products in themselves; at present they are among the few amino acids that are not predominantly industrially produced by bacterial fermentation (Wendisch, 2019).

In yeast, AAA synthesis begins with the shikimate pathway (Fig. 1) wherein the DAHP synthases Aro3p and Aro4p convert erythrose-4-phosphate (E4P) and phosphoenolpyruvate (PEP) to 3-deoxy-D-arabinoheptulosonate-7-phosphate (DAHP). This is in turn converted to chorismate via Aro1p and Aro2p. Chorismate is then either committed to phenylalanine/tyrosine biosynthesis via conversion to prephenate by Aro7p or tryptophan biosynthesis via conversion to anthranilate by Trp2p/3p. In the former case, the enzymes Pha2p or Tyr1p then convert prephenate to phenyl- or hydroxyphenylpyruvate, which are then transaminated to phenylalanine or tyrosine respectively by Aro8p and Aro9p. In native biosynthetic pathways, Aro3/4p and Aro7p are feedback inhibited by tyrosine and/or phenylalanine as committing to this pathway is metabolically expensive. To overcome this limitation, feedback-resistant alleles of all three genes have been identified and exploited to overproduce these AAAs. Typically, overexpressing feedback-resistant (fbr) *ARO4* and *ARO7* is sufficient to overproduce phenylalanine and tyrosine for the production of secondary metabolites (Koopman et al., 2012; Luttik et al., 2008), while an fbr allele of *ARO3* improves AAA production in combination with *ARO4* (Reifenrath et al., 2018a). Beyond this, various studies have partially or completely overexpressed the shikimate or phenylalanine/tyrosine pathways to further direct flux down this pathway. For example, E4P supply to the pathway can be increased in *S. cerevisiae* by partially overexpressing the non-oxidative branch of its pentose phosphate pathway (PPP) (Curran et al., 2013; Suástegui et al., 2017) and PEP supply to the shikimate pathway has been increased by increasing its natural production or slowing its conversion to pyruvate (Hassing et al., 2019; Mao et al., 2017).

**Figure 1.**
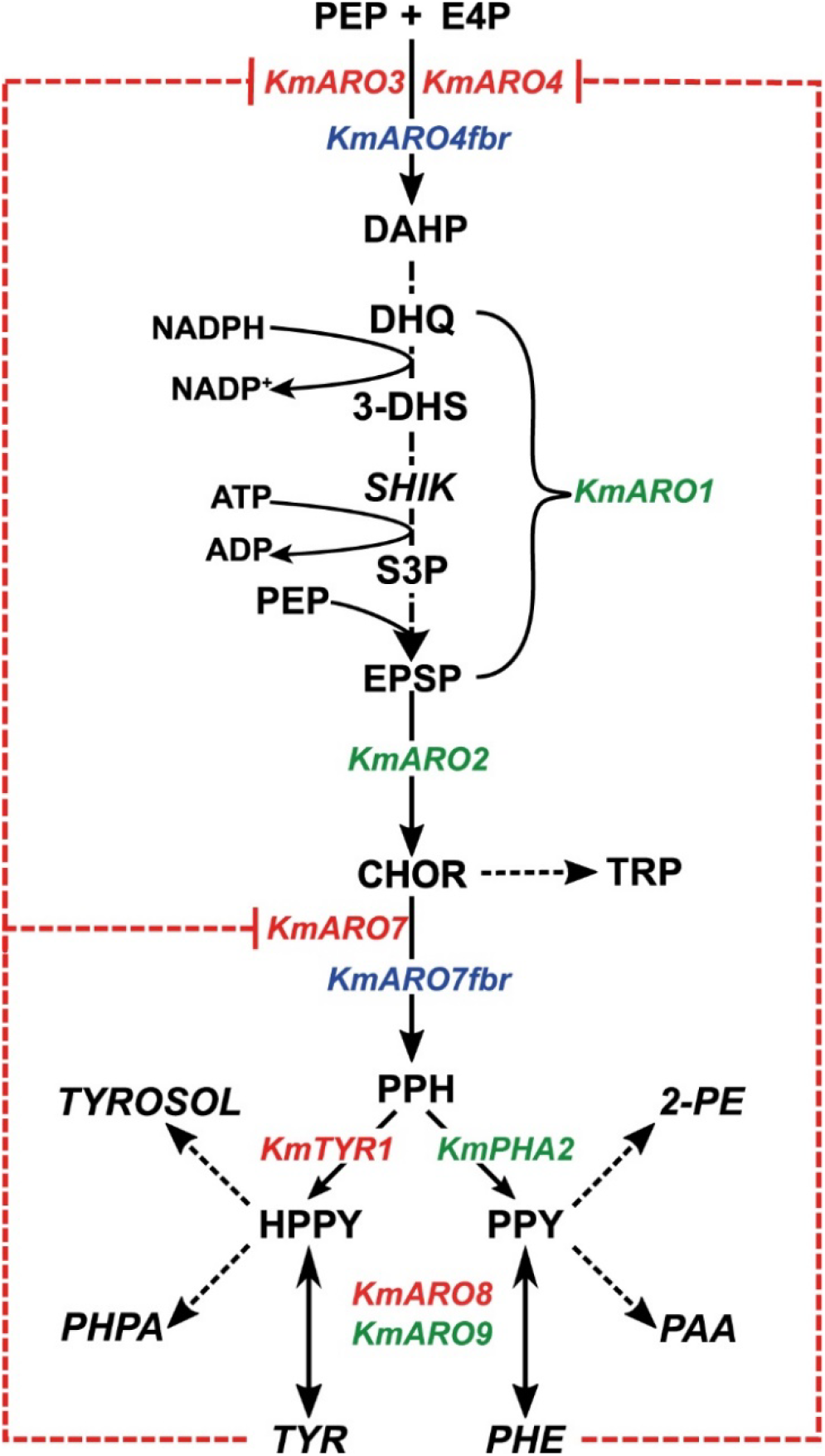
Shikimate and phenylalanine/tyrosine biosynthesis in *Kluyeromyces marxianus*. The genes are named according to their orthologues in *Saccharomyces cerevisiae*. Genes in red are deleted or otherwise inactivated, genes in green are overexpressed in our study, and genes in blue are overexpressed alleles of genes resistant to feedback inhibition. Red dashed lines indicate product inhibition, while dashed arrows indicated multiple reactions between the metabolites indicated. Compounds that were assayed by HPLC are italicised. Abbreviations: PEP: phosphoenolpyruvate, E4P: erythrose 4-phosphate, DAHP: 3-deoxy-D-arabinoheptulosonate-7-phosphate, DHQ: 3-dehydroquinate, 3-DHS: 3-dehydroshikimate, SHIK: shikimic acid, S3P: shikimate-3-phosphate EPSP: 5-enoylpyruvateshikimate-3-phosphate, CHOR: chorismate, PPH; prephentate, (H)PPY: (hydroxyl)phenylpyruvate, 2-PE: 2-phenylethanol; PAA: phenylacetic acid; TYROSOL: tyrosol/para-hydroxyphenylethanol; PHPA: para-hydroxphenylacetate; TRP: tryptophan; TYR: tyrosine, PHE: phenylalanine.

While *S. cerevisiae* remains a favoured host, for several reasons, there is considerable value in exploring alternative yeast platforms for the synthesis of aromatic compounds. First, the native metabolism of *S. cerevisiae* has evolved for ethanol production, often at the expense of other pathways. Thus, the metabolism of a Crabtree-negative yeast can offer a better starting point since carbon flux is not diverted to ethanol and the carbon distribution between the various pathways that give rise to anabolic precursors is better balanced. Second, some non-S*accharomyces* yeasts possess advantageous traits such as rapid growth, thermotolerance, salt tolerance or the capacity to use a wider array of substrates (Radecka et al., 2015). The development of alternative hosts has previously been delayed by an insufficient knowledge of cellular physiology, limited genetic tools, and a lack of experience of developing scaled bioprocesses. The rapid growth in the implementation of omics technologies, coupled with the development of synthetic biology (Löbs et al., 2017), is helping address the first two of these problems and several non-*Saccharomyces* yeasts are close to crossing the threshold to becoming credible first-choice hosts for defined applications.

One such yeast is the Crabtree-negative *Kluyveromyces marxianus*. It can tolerate temperatures over 50°C and has the fastest growth rate of any eukaryote (Gombert et al., 2016; Lane and Morrissey, 2010). It can natively consume pentose sugars and disaccharides found in agricultural and dairy wastes, making these economic feedstocks for potential cell factories. The past decade has seen development of tools for its metabolic engineering such as CRISPR/Cas9 systems, libraries of regulatory elements, secretory peptides, and *in vivo* assembly protocols (Nurcholis et al., 2020). In parallel, *K. marxianus* has been successfully used as a platform to produce heterologous compounds such as short-chain alcohols, carotenoids and polyketides (Cheon et al., 2014; Lin et al., 2017; McTaggart et al., 2019). However, all the latter studies focus on the overexpression of heterologous enzymes or pathways and only a few studies attempt engineer metabolism to improve production (Cernak et al., 2018; Kong et al., 2019). With regards to aromatic compounds, *K. marxianus* has been evolved to overproduce phenylalanine and by extension the aromatic alcohol 2-phenylethanol (2-PE) via an enhanced Ehrlich pathway (Kim et al., 2014), but this has shed little light on appropriate metabolic engineering strategies. As its physiology relies more on respiration than *S. cerevisiae*, approaches to metabolically engineer *K. marxianus* to maximise precursor supply and product titres and yields can vary significantly from known ones in baker’s yeast, especially since the production of aromatics is not a fermentative process.

In this work, we adopt a structured approach to metabolically engineer *Kluyveromyces marxianus’* AAA biosynthetic pathway to overproduce phenylalanine. Our strategy follows three stages. First, we alleviate feedback inhibition of the native shikimate pathway to establish AAA overproduction, followed by overexpression of the phenylalanine biosynthetic enzymes to enhance the production of phenylalanine. Second, we attempt to increase the supply of E4P and PEP to the shikimate pathway, first independently and then simultaneously, using native and heterologous enzymes. Finally, we redirect flux to phenylalanine synthesis alone by knocking down competing synthetic and degradative pathways. These strategies were assessed individually and in combination, and multiple different strategies aimed at similar goals were also evaluated. Following this strategy allowed us to increase and redirect flux to the phenylalanine pathway so that it comprises up to 90% of the aromatic products synthesized. In the course of our work, we track the production 2-phenylethanol (2-PE), the Ehrlich degradation alcohol of phenylalanine as a proxy for the production of this amino acids. Our systematic investigation of engineering strategies for phenylalanine overproduction provide a roadmap for future metabolic engineers and synthetic biologists wishing to work with *Kluyveromyces marxianus*.

## 2. Materials and methods

### 2.1. Strains and media

*Kluyveromyces marxianus* strain NBRC1777 from the Biological Resource Centre, NITE (NBRC; Tokyo Japan) was used as the base strain for all work in this study (Inokuma et al., 2015). For preliminary genome editing and knock-outs, strains were routinely grown in YPD broth (2% yeast extract, 1% peptone and 2% glucose; Formedium, Hunstanton, UK), or synthetic drop-out (SD) medium with the relevant supplements (0.5% ammonium sulphate, 0.19% yeast nitrogen base without amino acids or ammonia (Formedium), 2% glucose and the relevant drop-out supplement according to the manufacturer’s recommendations (Formedium)). When cells needed to be grown on solid media, the recipes above were supplemented with 2% agar (Formedium). For transformations involving CRISPR/Cas9, strains were grown on YPD agar with 200 mg L^−1^ hygromycin for selection. Bacterial transformations used *E.coli* DH5a grown in LB medium (1% NaCl, 1% peptone, 0.5% yeast extract; Formedium) or LB agar (+1.5% agar) with the appropriate antibiotic (100mg L^−1^ ampicillin, 50mg L^−1^ chloramphenicol or 50mg L^−1^ kanamycin) as required. All antibiotics and reagents were purchased from Fisher Scientific (Dublin, Ireland) or Sigma-Alrdich/Merck (Haverhill, UK). All primers were obtained from Sigma-Aldrich.

### 2.2. Plasmid construction

All genes for the AAA biosynthetic pathway and non-oxidative PPP were amplified from genomic DNA from *K. marxianus* strain CBS6556 (Westerdijk Fungal Biodiversity Institute, Utrecht, The Netherlands) using Q5 High-Fidelity Polymerase (M4092L, New England Biolabs (NEB), Ipswich, UK) with flanking nested BsaI (R0535L, NEB) and BsmBI (R0739L, NEB) restriction enzyme sites added by PCR. All synthetic heterologous genes were codon-optimised for *S. cerevisiae* and obtained from Twist Biosciences (San Francisco, U.S.A.).These respectively were used to clone the genes into storage plasmids (level I) by Golden Gate assembly with BsmBI and T7 DNA ligase (M0318L, NEB) for subsequent assembly into expression cassette-bearing plasmids (level II) using the hierarchical system based on the Yeast Toolkit standard (Lee et al., 2015). Component vectors containing markers and connectors used to construct cloning vectors using this standard were a gift from John Dueber distributed through (Kit # 1000000061, Addgene, Watertown, U.S.A.). The genes or gene fragments were cloned into the storage vector pYTK001 using a Golden Gate assembly with BsmBI. This plasmid contains a GFP cassette in its ‘empty’ state, allowing for green/white colony screening. Each plasmid’s insert was verified by colony PCR following transformation into *E.coli,* and subsequent sequencing of the inserts of PCR-positive plasmids. For constructing feedback-resistant alleles of *KmARO4* and *KmARO7*, the mutations were made to the genes during Golden Gate assembly by amplifying the gene as two fragments flanking the residue to be mutated. The mutated residue (K221L in *KmARO4* and G141S in *KmARO7*) was included in the complementary overhangs for the BsmBI enzyme sites meant to join the fragments, so that on successful assembly of the fragments into a level I plasmid, the cloned product is the entire gene with the incorporated mutation. A similar strategy was used to eliminate internal BsaI or BsmBI sites that would otherwise interfere with Golden Gate assembly.

Level I plasmids for each gene and the desired promoter and terminator were then assembled into transcriptional units (TUs) by Golden Gate assembly with BsaI/T7 DNA ligase, which were then transformed into *E.coli* and verified by colony PCR using OneTaq Quick-Load Polymerase (M0486L, NEB) and restriction digestion by NotI (FD0596, Fisher Scientific). The *K. marxianus* promoters and terminators used for these plasmids have been described elsewhere (Rajkumar et al., 2019).Depending on their intended positions in a multi-TU plasmid, TUs in level II plasmids were cloned with direction-specific connectors flanked by BsmBI sites provided in the Yeast Toolkit to ensure directional assembly. Depending on the planned constructs, some TUs were cloned multiple times, with different connectors. Finally, multiple TUs comprising pathways or parts of pathways were cloned into multi-TU plasmids by a Golden Gate assembly of the TU plasmids into drop-out expression vectors constructed in-house using BsmBI/T7 DNA ligase. All the pathway expression plasmids and cloning vectors are listed in Table 1. All level I and level II plasmids constructed for this study are listed in Table S1, and the primers used to create them in Table S2. More detailed information on Golden Gate assembly of the plasmids can be found in refs (Lee et al., 2015; Rajkumar et al., 2019).

**Table 1.**
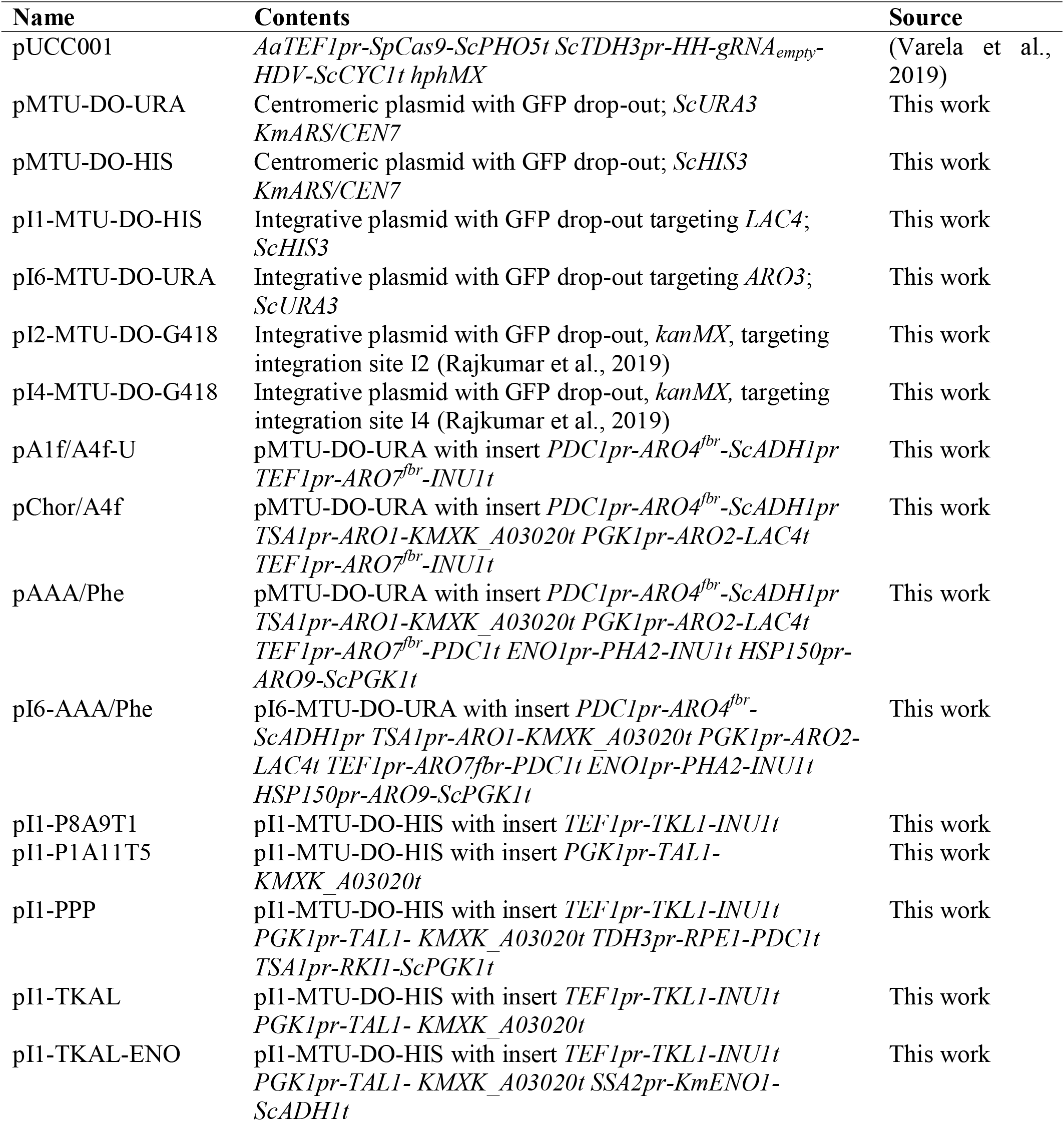
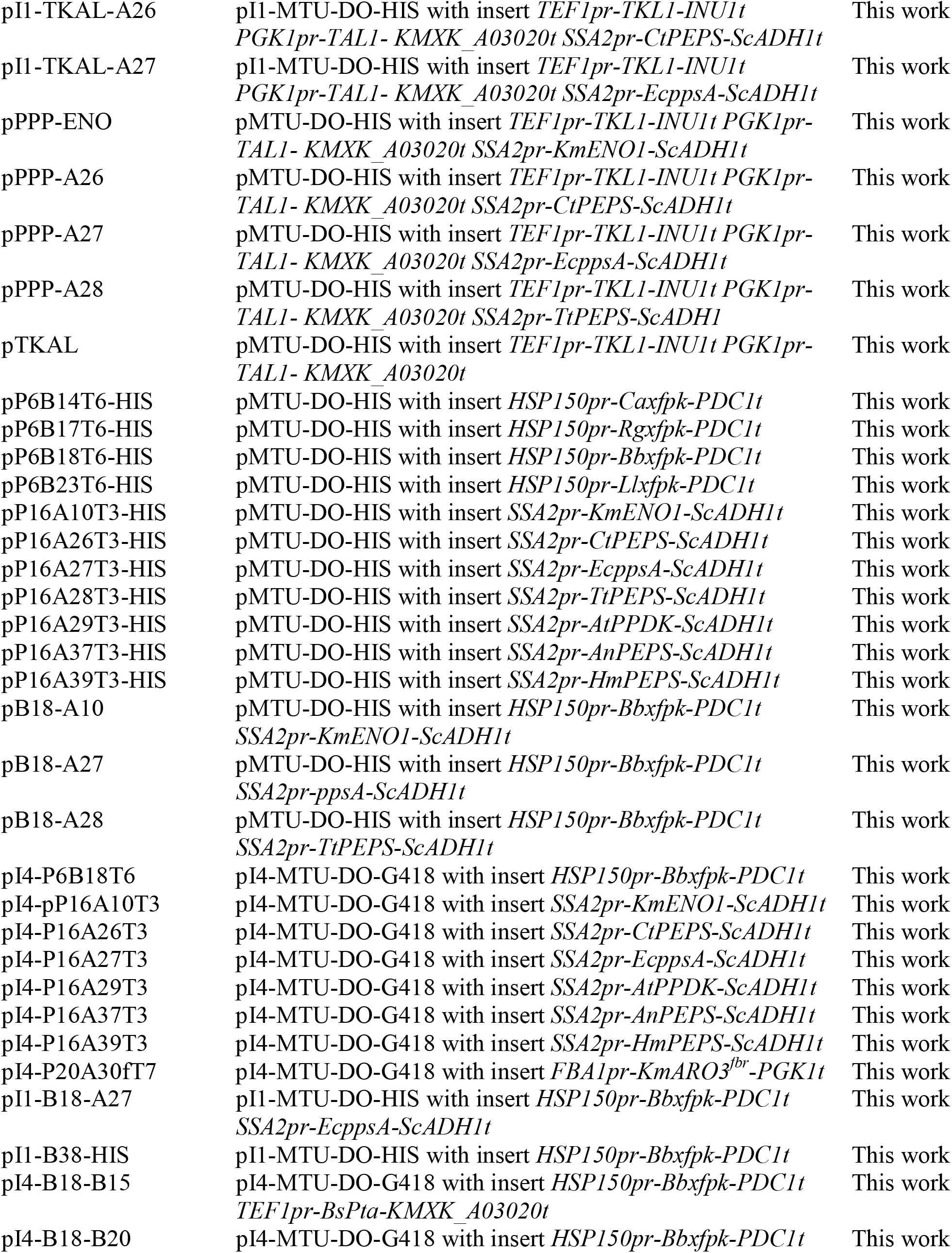
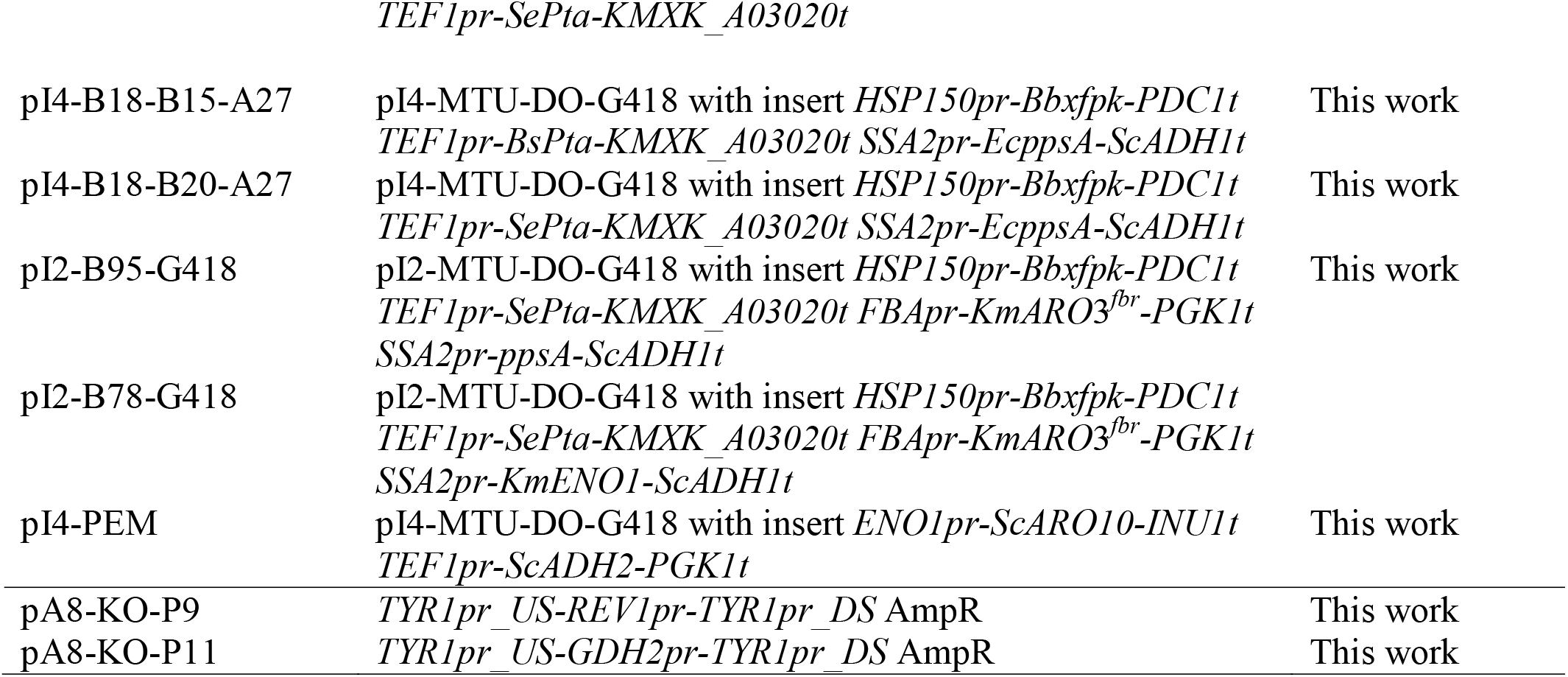
Expression plasmids used in this study. Unless specified otherwise, all genes, promoters and terminators in the inserts are from *K. marxianus.*

### 2.3. Production strain construction

#### 2.3.1. Strain background engineering

For plasmid-based experiments evaluating the overexpression of *KmARO4*^*fbr*^, *KmARO7*^*fbr*^, and the shikimate and phenylalanine pathways, we used the ura-auxotrophic strain KmASR.006 as a background. To create a background strain missing wild-type *ARO3, ARO4* and *ARO7* we sequentially inactivated the genes using our CRISPR/Cas9 system pUCC001, which contains a hygromycin resistance marker. As previously described, gRNAs targeting these genes were designed using sgRNA and cloned into pUCC001 by Golden Gate assembly (Varela et al., 2019; Xie et al., 2014). Starting with a ura^−^ his^−^ auxotroph KmASR.008 we had previously constructed, we first inactivated *KmARO4* and then *KmARO7* sequentially. In each case the CRISPR plasmid with the relevant gRNA target was transformed into KmASR.008 and selected for on YPD containing hygromycin. Up to 8 colonies were screened for mutations at *KmARO4* or *KmARO7* that caused a shift in the reading frame. The deletions provided us with strain KmASR.029 with an *aro4 aro7* genotype. *KmDNL4* was subsequently inactivated in this strain by CRISPR in a similar fashion to yield KmASR.039 to allow specific integration of pathway vectors via inactivated non-homologous end-joining (NHEJ)(Nambu-Nishida et al., 2017). After each successful gene inactivation, the strains were cured of the CRISPR plasmid by passaging an overnight culture of the confirmed mutant strain twice into YPD before being used for further genome editing. Mutant strains had their targeted genes sequenced again following curing to ensure the mutation was stable. All strains used and constructed in this study are listed in Table 2, and all gRNA sequences used to inactivate genes are listed in Table S3.

**Table 2.**
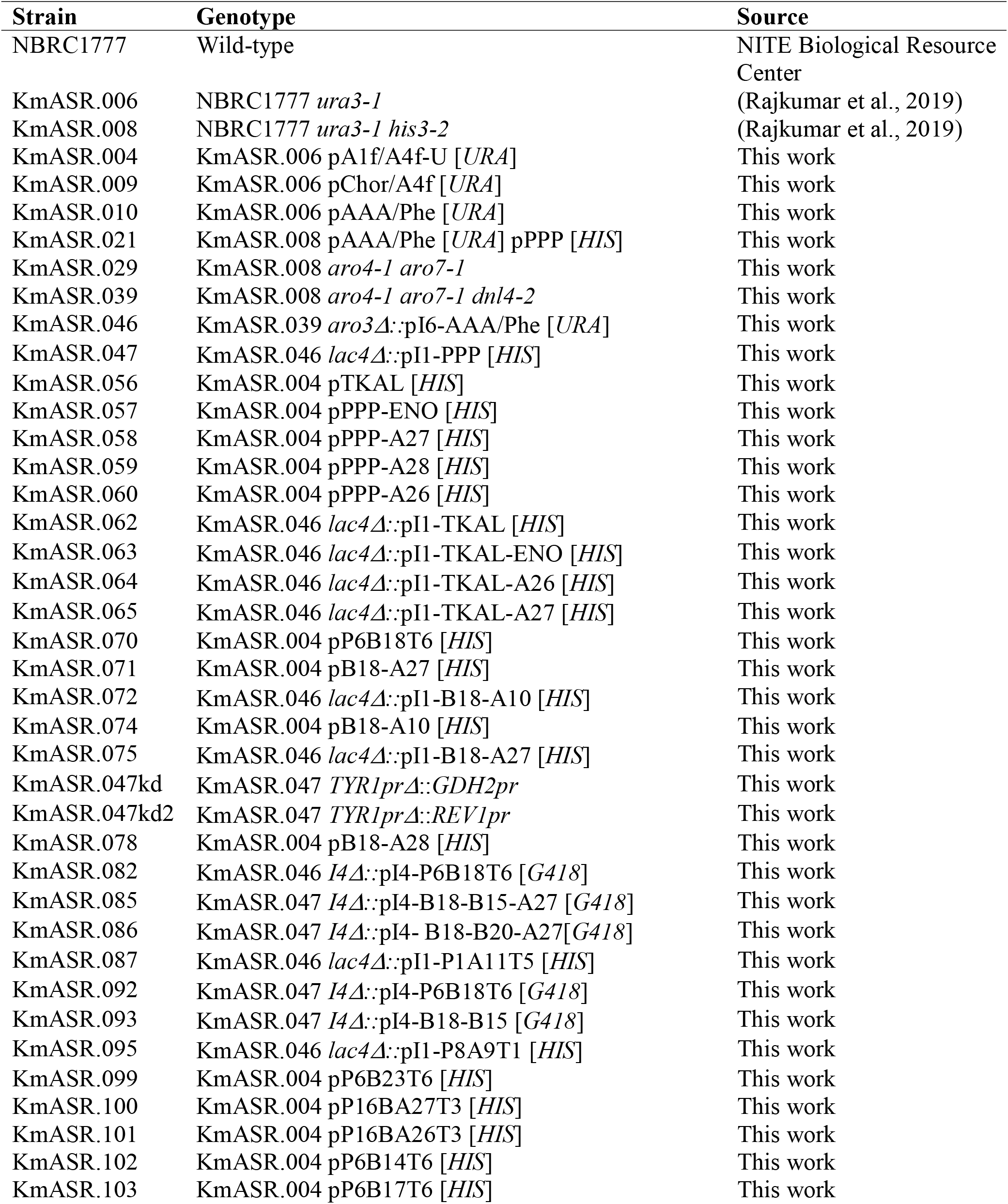
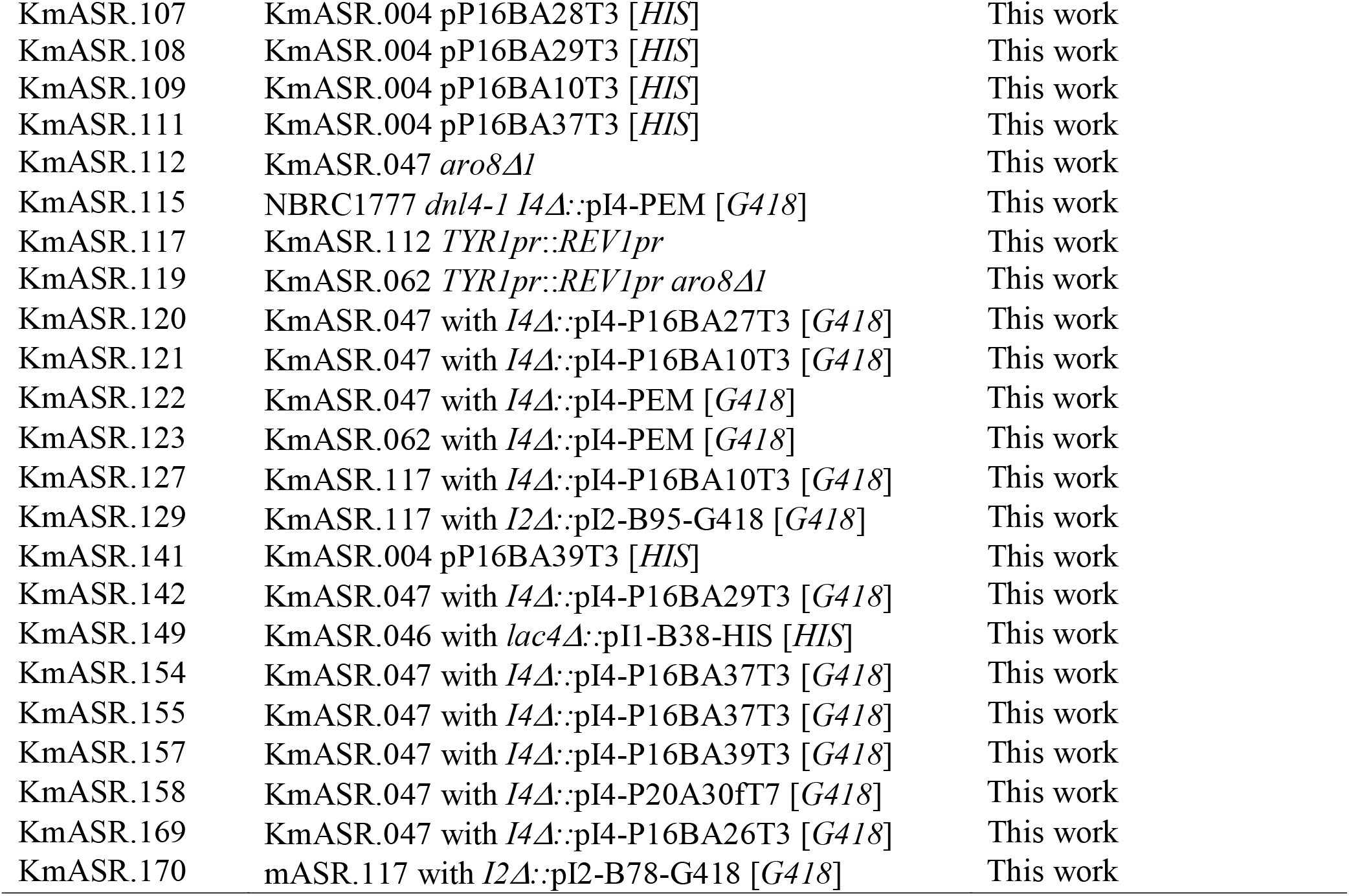
Strains used in this study.

#### 2.3.2 Transformations and strain confirmation

All transformations into *K. marxianus* were carried out by the lithium acetate/PEG method (Gietz, 2014). The expression vectors were either centromeric or integrative and were constructed as recommended for the Yeast Toolkit standard using a GFP drop-out cassette whose loss confirms successful assembly and allows for green/white colony screening of *E.coli* transformed with the Golden Gate assembly. All cloning vectors used in this work have been deposited in Addgene, with their part numbers listed in Table S5. For strains establishing the overexpression of the feedback-resistant shikimate or phenylalanine pathway, plasmids of interest were transformed into KmASR.008 and selected for on SD-ura agar. Subsequent plasmids containing genes to increase precursor supply (Fig. S1) were transformed in the resultant strains and selected for on SD-ura-his agar. For the strain with the integrated feedback-resistant phenylalanine pathway KmASR.046, the pathway was cloned into the integrative vector pI6-MTU-DO-URA targeting *KmARO3*, resulting in the vectors pI6-AAA/Phe. 1µg of this plasmid was digested with FastDigest SgsI (FD1894, Fisher Scientific), transformed into KmASR.039 and selected on SD-ura agar. Colonies appearing 3-5 days after transformation were screened by duplex colony PCR: one pair of primers checking integration of the pathway, priming upstream of the *ARO3* locus and at the 5’ end of the construct, and another pair priming downstream of the *ARO3* locus and at the 3’ end of the construct. KmASR.046 was selected from colonies positive for both bands. For the overexpression of genes increasing precursor supply in KmASR.046, genes of the non-oxidative pentose phosphate pathway or phosphoketolases/phosphotransacetlyase were cloned into the vector pI1-MTU-DO-HIS with a histidine auxotrophic marker targeting the *LAC4* locus and KmASR.047 respectively. Correct integration of the relevant construct was verified by colony PCR using two pairs of primers designed in a manner similar to those for checking the integration of pI6-AAA/Phe. The subsequent overexpression of genes in the resulting strains (Table 2, Fig. S1) were done using the cloning vectors pI2-MTU-DO-G418 and pI3-MTU-DO-G418 that target integration sites on chromosomes V and IV respectively and previously characterized by us (Rajkumar et al., 2019). Primer sequences for colony PCR verification are listed in Table S2.

#### 2.3.3. Tyrosine knockdown by CRISPR/Cas9

For marker-free replacement of *TYR1pr*, a gRNA target for Cas9 covering the first 13 bases of *KmTYR1* and the first 7 upstream bases was identified by examining the start of the gene and its 5’ UTR. These were used to construct gRNA plasmid pJ22. The *REV1* or *GDH2* promoters were cloned by Golden Gate assembly into a marker-free integrative vector flanked by 850bp homology arms directly upstream and downstream of the gRNA sequence (while making sure the knock-down promoter was in-frame of *KmTYR1*) to yield plasmids pA8-KO-P9 and pA8-KO-P11 respectively. These plasmids were linearized with NotI and transformed into KmASR.047 alongside pJ22. Hygromycin-resistant colonies were screened for promoter replacement by colony PCR using a forward primer priming in the knock-down primers and a reverse primer at the end of *KmTYR1*.

#### 2.3.4. Gene deletions

gRNA targets for *KmARO8* were identified and cloned into pUCC001 as previously described. However, as these were to be transformed into an NHEJ-deficient background, it was necessary to provide a linear repair fragment for homology-directed repair. These were designed by fusing two 80bp sequences flanking a portion of the gene to be deleted (440-650bp; the size of the deletion depends on the position of the target and the colony PCR assay). The repair fragment was ordered as two 90bp oligos with a 20bp region in common (the last 10bp of each flanking sequence). 50pmol of each primer was then annealed and extended by Q5 polymerase in a short PCR reaction without a template to create a 160bp repair fragment, which was in turn transformed with the relevant gRNA plasmid targeting *KmARO8* (pJA6). Hygromycin-resistant colonies were screened by colony PCR for the deletion by primers amplifying the entire gene; the difference in PCR product size relative to an intact gene indicated a successful deletion. Ultimately *KmARO8* was knocked out in KmASR.047 to yield KmASR.112, and the deletion repeated out in the *TYR1* knock-down strain ASR.047kd2 to create KmASR.117.

### 2.5. Production strain cultivation and sampling

The strain whose production was to be studied was grown overnight in 5mL of the SD medium (Formedium) at 30°C in a shaking incubator set to 200rpm. This culture was then diluted to an OD of 0.1 in 20mL of minimal medium (5g L^−1^ (NH_4_)_2_SO_4_, 3g L^−1^ KH_2_PO_4_, 0.5g L^−1^ MgSO_4_.7H_2_O, 20 g L^−1^ glucose with vitamins and trace elements, pH 6) (Fonseca et al., 2007) in a 100mL conical flask and grown at 30°C and 200rpm. 1mL samples were taken at 48h to sample extracellular metabolites. When more time points were sampled, a smaller volume of the culture medium was taken at each time point. After the OD of the culture was measured, the culture medium was centrifuged at 13,000rpm for 10 minutes and the supernatant recovered and stored at −20°C until analysis. For the measurement of intracellular metabolites a pellet from 2mL culture was boiled with 75% ethanol (Lai et al., 2017), the supernatant recovered and stored at −20°C until analysis.

### 2.6. Analytical methods

Extra- and intra-cellular metabolite samples were thawed, vortexed, and transferred to screw-cap HPLC vials at an appropriate dilution. Shikimic acid, phenylalanine, tyrosine, 2-phenylethanol, tyrosol and para-hydroxyphenylacetic acid were analysed on an Agilent 1260 Infinity HPLC Infinity system with a Zorbax Eclipse Plus C18 column (4.6 × 150mm, 3.5µm; Agilent, Little Island, Ireland) operating at 40°C using 20mM KH_2_PO_4_ with 1% acetonitrile (pH 2)/acetonitrile as a solvent system at a flow rate of 0.8 mL min^−1^ as previously described (Koopman et al., 2012). The elution gradient was as follows: acetonitrile was increased from 0-10% for the first 6 minutes, then increased to 40% between 6 to 23 minutes, and then switched to 100% KH_2_PO_4_ for 23 to 27 minutes. All metabolites were detected using a VWD100 multi-wavelength detector measuring the absorbance at 200nm. Data were analysed using OpenLab CDS (Agilent). The elution times of the metabolites of interest were as follows: shikimic acid, 2.32 minutes; tyrosine, 5.4 minutes; phenylalanine, 7.5 minutes; tyrosol/para-hydroxyphenylethanol, 10.22 minutes; para-hydroxyphenylacetic acid, 11.17 minutes; 2-phenylethanol, 17.2 minutes; phenylacetic acid; 17.86 minutes. Standards for shikimic acid, phenylalanine, tryptophan, 2-phenylethanol and phenylacetic acid were obtained from Sigma-Aldrich, whereas standards for tyrosol and para-hydroxyphenylacetic acid were obtained from Fisher Scientific. Extracellular acetate concentration was determined with an Agilent 1200 HPLC system (Agilent Technologies, CA, USA) with a REZEX 8 μL 8% H+ organic acid column (300 ×7.8 mm) (Phenomenex, CA, USA) operating at 65◦C using 0.01 N H_2_SO_4_ as a solvent at a flow rate of 0.6 mL min^−1^. Organic compounds were detected with a refractive index detector.

## 3. Results

### 3.1. Establishing feedback-resistant phenylalanine biosynthesis in *Kluyveromyces marxianus*

The aromatic amino acid biosynthetic and degradative pathways in *K. marxianus* share all steps in common with *S. cerevisiae* (Fig. 1, Table S4) though there are some differences in numbers of paralogous genes. While *S. cerevisiae* has three phenylpyruvate decarboxylases Pdc5p, Pdc6p and Aro10p, *K. marxianus* only has a homologue to *ARO10* and a putative one to *PDC5* whose enzymatic activity is unknown (Choo et al., 2018). In contrast, besides homologues of the two aminotransferases found in *S. cerevisiae* (*ARO8* and *ARO9*), sequenced *K. marxianus* genomes contain a third gene annotated as a putative aminotransferase (Lertwattanasakul et al., 2015). The rational synthetic biology approach that we adopted involved the construction and analysis of over 60 strains (Table 2) in a complex matrix as we tested ways to enhance and optimise flux through the AAA biosynthetic pathway. A chart that shows this matrix and the relationship of the different strains is provided to aid understanding (Fig. S1).

We began our engineering strategy by alleviating feedback inhibition of AAA synthesis in *K. marxianus*, a key step in establishing phenylalanine or tyrosine overproduction in yeast (Luttik et al., 2008). The *K. marxianus* enzymes (KmAro4 and KmAro7) share >75% sequence identity with their *S. cerevisiae* orthologues (Table S4), thus we used knowledge of *S. cerevisiae* feedback resistant mutations as the basis of generating feedback resistant variants in *K. marxianus* enzymes. The substitutions K221L in Aro4^fbr^ and G141S in KmAro7^fb*r*^ are equivalent to K229L and G141S mutations in the same enzymes in *S. cerevisiae* (Luttik et al., 2008). We initially overexpressed single copies of *KmARO4*^*fbr*^ and *KmARO7*^*fbr*^ together in *K. marxianus* strain NBRC1777 on the centromeric plasmid pA1f/A4f-U. The resultant strain, KmASR.004, produced detectable amounts of 2-PE and tyrosol (the Ehrlich alcohol derived from tyrosine), in the culture medium as a result of AAA overflow from the feedback resistant enzymes, along with significant amounts of shikimate. Approximately 1.2-1.4 mM, or 170 mg L^−1^ of each aromatic alcohol was produced after 48h shake-flask culture in minimal medium at 30°C along with 1.6mM shikimate, whereas the wild-type yeast did not produce any of these metabolites when grown for the same length of time (Fig. 2). This established that feedback resistant variants can be used in *K. marxianus* to alleviate feedback inhibition by both tyrosine and phenylalanine.

**Figure 2.**
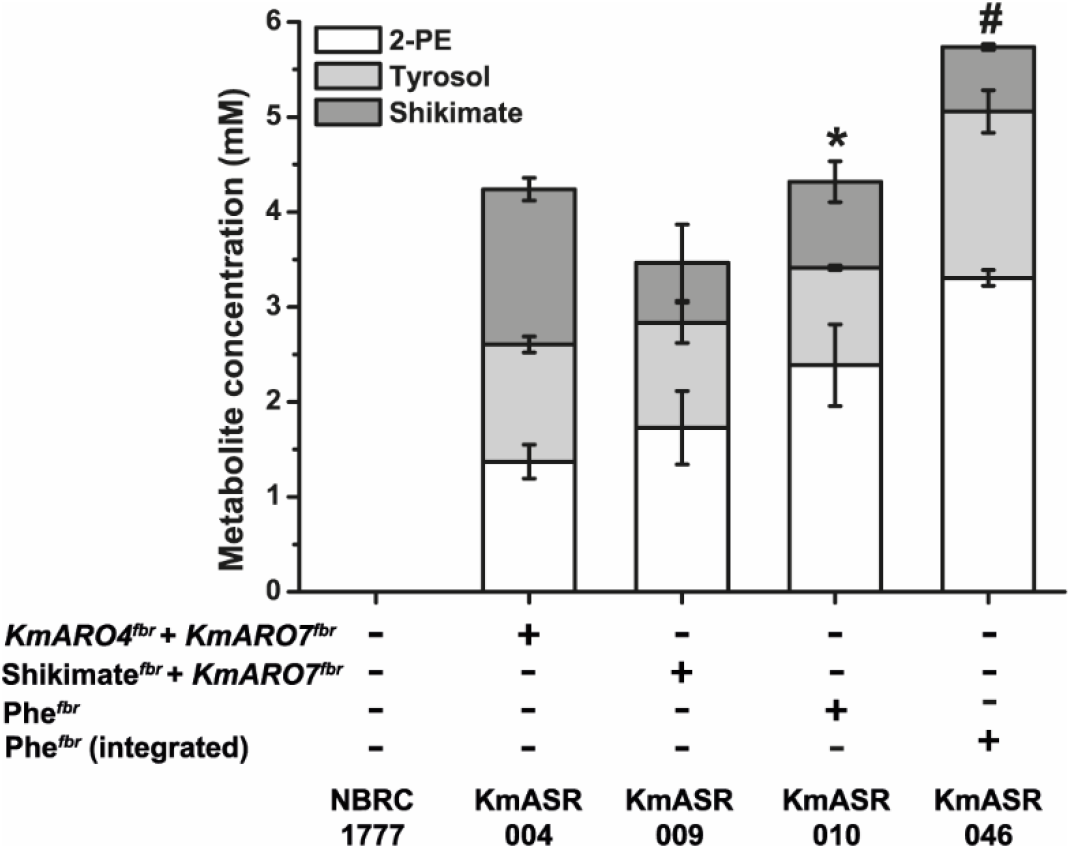
Overexpression of a feedback-resistant phenylalanine biosynthetic pathway. The two feedback-resistant alleles of *KmARO4* and *KmARO7* constructed based on homology to *S. cerevisiae* are functional, with their episomal overexpression leading to the production of Ehrlich alcohols of phenylalanine (white) and tyrosine (grey) from the overflow of their increased synthesis, as well as excess shikimate (dark grey). All compounds were measured by HPLC in culture medium after 48h growth at 30°C as described in the Methods. Subsequent co-expression of the shikimate (*KmARO4*^*fbr*^, *KmARO1, KmARO2*) + *KmARO7*^*fbr*^ and phenylalanine (*KmARO4*^*fbr*^ + KmARO1 + KmARO2 + *KmARO7*^*fbr*^ +*KmPHA2*+*KmARO9*) pathways increased their production and converted more shikimate to amino acids. Chromosomal integration of the pathway further improved phenylalanine and tyrosine production. Strains marked with an asterisk (*) show a significantly higher level of 2-PE relative to that produced by ASR.004 as determined by a paired *t-*test (*p*<0.05). The data in the bar plots are the mean ± s.d. of at least three replicates. All data plotted in this and following graphs can be found in Supplementary Data 1.

The next step was to create the centromeric plasmid pChor/A4f overexpressing both feedback resistant alleles (*ARO4*^*fbr*^ and *ARO7*^*fbr*^) and two other genes of the shikimate pathway, *KmARO1* and *KmARO2.* The KmASR.009 strain bearing this plasmid showed an increase in pathway flux with an increase of 25% in 2-PE titres relative to KmASR.004 (Fig. 2) as well as reduced levels of the intermediate shikimate. To further push flux coming from chorismate to phenylalanine, we further overexpressed *KmPHA2* and *KmARO9* in a new single plasmid pAAA/Phe. The reason that *KmARO9* is overexpressed is to facilitate our approach of using 2-PE as a readout for flux towards phenylalanine (Fig. 1). Strain KmASR.010 with these additions increased 2-PE titres by 40% relative to KmASR.009, ultimately comprising nearly 70% of the measured Ehrlich metabolites (Fig.2). The increased pull of the remaining enzymes downstream of Aro7p led to less shikimate being secreted but instead further utilized downstream, with the acid only comprising 20% of the total aromatic products as opposed to nearly 40% for ASR.004. At this stage, it appeared that all excess phenylalanine or tyrosine was being converted to 2-PE or tyrosol respectively; the amino acids were neither detected as extracellular nor intracellular metabolites. We also did not detect any intracellular Ehrlich metabolites, suggesting that they were all secreted during production.

We then decided to chromosomally integrate and overexpress the same pathway in a strain lacking the native alleles of *ARO3, ARO4* and *ARO7*. Besides more stable expression, such a background would ensure that *K. marxianus* relies solely on the feedback-resistant phenylalanine/tyrosine pathway. Using CRISPR/Cas9, we inactivated *KmARO4* and *KmARO7* in a non-homologous-end-joining (NHEJ)-deficient background. At the same time, we reconstructed the Phe^fbr^ pathway in an integrative vector pI6-AAA/Phe. We then integrated the pathway at the *KmARO3* locus to create strain KmASR.046, forcing *K. marxianus* to use this pathway for all AAA biosynthetic requirements. The combination of stable chromosomal integration of the pathway and loss of feedback-inhibited genes resulted in a final increase of nearly 40% in 2-PE production in the resultant strain KmASR.046 compared to plasmid-based expression in KmASR.010, with a final titre of 400 mg L^−1^ 2-PE and 241 mg L^−1^ tyrosol (3.31mM and 1.75mM, Fig. 2).

### 3.2. Increasing precursor supply to the shikimate pathway

#### 3.2.1. Increasing phosphoenolpyruvate supply to the shikimate pathway

We next decided to investigate the best means to increase phosphoenolpyruvate (PEP) and erythrose-4-phosphate (E4P) supply to the shikimate pathway. PEP is a substrate in two steps of the shikimate pathway, required by Aro4p and Aro1p (Fig. 1). In yeast, PEP is produced by enolase (Eno1/2p) during glycolysis and PEP carboxykinase (Pck1p) during gluconeogenesis (Fig. 3(A)). In other species, PEP is also synthesized by PEP synthases (PEPS, EC 2.7.9.2) or pyruvate, orthophosphate dikinase (PPDK, EC 2.7.9.1), enzymes which reversibly produce PEP from pyruvic acid (Chastain et al., 2011). In *E.coli,* the PEPS gene *EcppsA* has been frequently used to increase precursor supply to produce aromatic amino acid-derived compounds (Juminaga et al., 2012; Patnaik and Liao, 1994). While *EcppsA* was unsuccessfully tested for a similar role in one *K. marxianus* strain (Kim et al., 2014), we decided to test several of these enzymes in our *K. marxianus* background. We overexpressed five codon-optimized PEPSes: *EcppsA*, an archaeal PEPS and three fungal PEPSes (Table 3). The fungal PEPSes were chosen based on their sequence homology to *EcppsA* while the archaeal PEPS was chosen since it was found to solely catalyse PEP formation (Chen et al., 2019). We also chose to test a codon-optimised PPDK from *Arabidopsis thaliana* as it has a 10-fold higher K_m_ for pyruvate than PEP as substrate, which could allow for a fastidious conversion of pyruvate to PEP (Chastain et al., 2011). We also included the native *ENO1* from *K. marxianus*. Each of these genes was individually introduced into ASR.004 and the effect on 2-PE production assessed (Fig 3B). All three classes of enzyme were able to increase aromatic amino acid and shikimate production providing more PEP for AAA synthesis. Among heterologous enzymes, the PEPS from *Aspergillus nomius* (ASR.111) and PPDK from *A. thaliana* (ASR.108) were the best performing, both increasing each 2-PE production to over 2mM (Fig. 3(B)). Overexpressing *KmENO1* proved to be the most effective strategy to increase PEP, producing nearly 2.4mM 2-PE, which represented an increase by over 70% relative to ASR.004 and nearly identical to the titre produced using the Phe^*fbr*^ (ASR.109, Fig. 3(B)).

**Figure 3.**
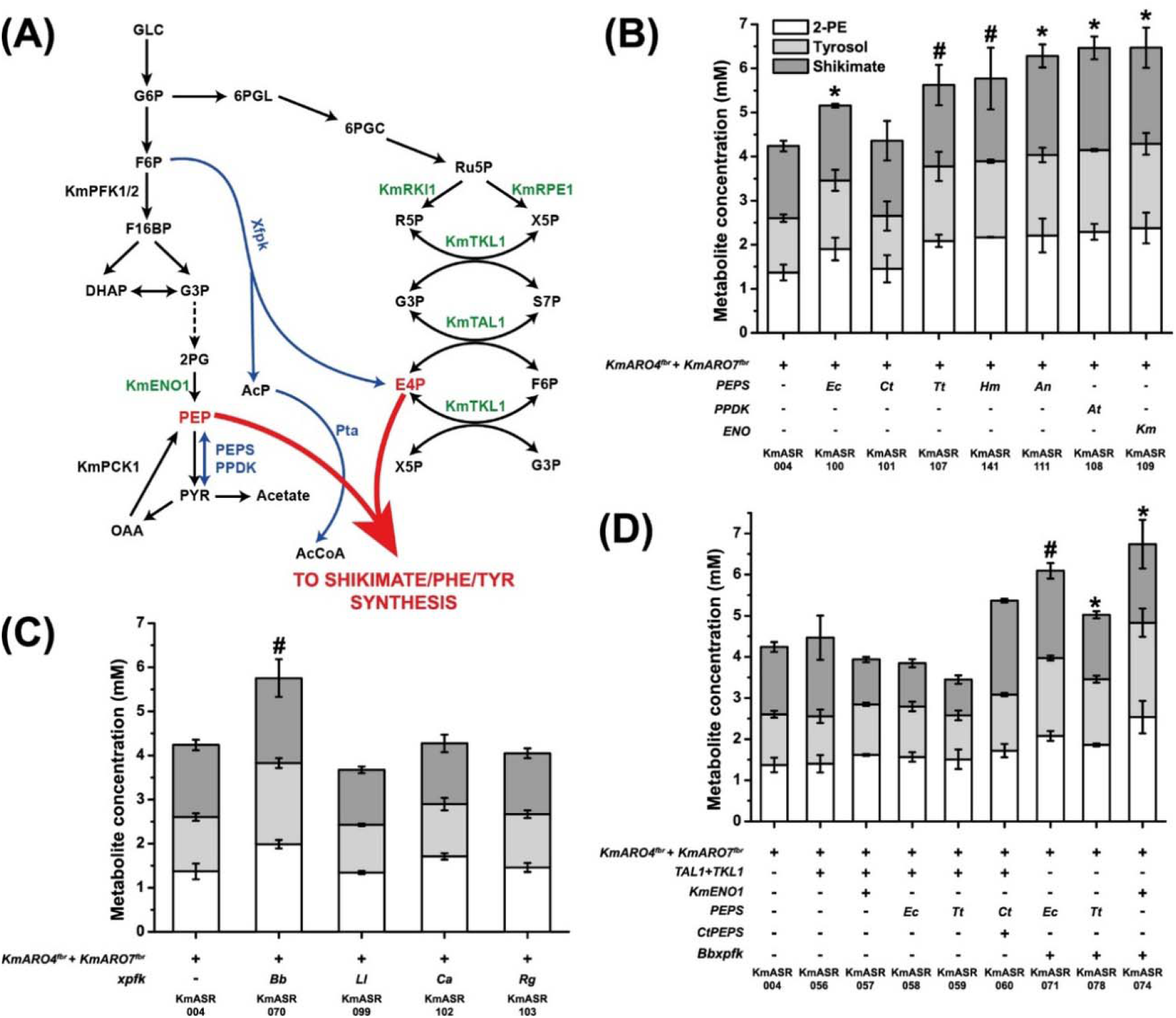
Engineering precursor supply to the shikimate pathway. **(A)** Overall view of the metabolic pathways giving rise to phosphoenolpyruvate (PEP) and erythrose-4-phosphate (E4P, and also the reactions that consume them. Reactions and genes in blue are heterologous, enzymes in green have their genes overexpressed, and other genes of interest are named. **(B)** Increasing PEP supply to the shikimate pathway heterologous using PEP synthases and pyruvate orthophosphate dikinases, and a native enolase. Enzymes of all three classes significantly increased levels of 2-PE, our proxy for phenylalanine production. **(C)** Increasing E4P supply to the shikimate pathway using fructose-6-phosphate phosphoketolases (Xfpk) from bacterial and fungal sources. **(D)** Attempts to simultaneously increase PEP and E4P supply to the shikimate pathway showed that the effects of increasing either metabolite individually are not additive, though in the end a modification was found that increased 2-PE production beyond what was achieved by individual precursors. Strains producing significantly higher amounts of 2-PE relative to that produced by ASR.004 as determined by a paired *t-*test are marked by an asterisk (*, *p*<0.05) or a hash (#, *p*<0.005). All data are plotted as the mean ± s.d. of at least three replicates. Abbrevations for organisms are: *Ec*: *Escherichia coli*, *Ct*: *Chaetomium thermophilum*, *Tt*: *Thielavia terrestris, Hm: Halofax mediterranei, An: Aspergillus nomius, At: Arabidopsis thaliana, Km: Kluyveromyces marxianus, Bb: Bifidobacter breve, Ll: Lactococcus lactis, Ca: Clostridium acetobutylicum*, *Rg: Rhodotorula graminis*.

**Table 3.**
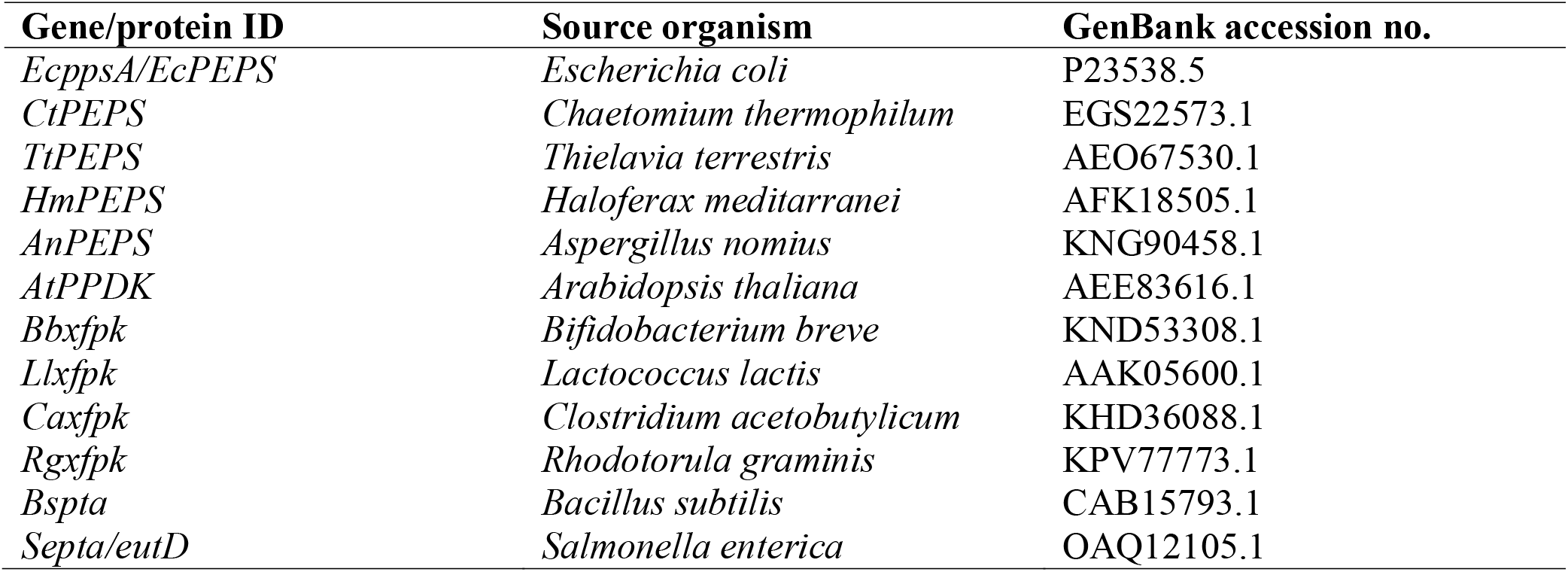
Gene name, source organism and GenBank ID for heterologous genes used in this study. Nucleotide sequences are provided in Supplementary Data 2.

#### 3.2.2. Increasing erythrose-4-phosphate supply to the shikimate pathway

The natural source for E4P is the non-oxidative branch of the pentose phosphate pathway (PPP) via cross-reactions from transketolase (Tkl1p) or transaldolase (Tal1p) (Fig. 3(A)). In *S. cerevisiae,* either or both of these enzymes are typically overexpressed to increase available E4P for AAA synthesis. More generally, metabolic engineering studies in *S. cerevisiae* have revealed that partial or complete overexpression of different parts of the non-oxidative PPP can increase the production of AAA or shikimate-derived products in combination with other modifications (Curran et al., 2013; Mao et al., 2017; Suástegui et al., 2017) As *K. marxianus* has previously been reported to have a higher flux through the PPP than *S. cerevisiae* (Blank et al., 2005; Jung et al., 2016), it was unclear whether overexpressing these enzymes would significantly improve E4P levels in *K. marxianus*. An alternative strategy tested in *S. cerevisiae* was to express a non-native fructose-6-phosphate phosphoketolase (Xfpk, EC 4.1.2.22, which converts fructose-6-phosphate to E4P and acetyl phosphate (Fig. 3(A)) (Bergman et al., 2019). We tested four active phosphoketolases reported in the literature: three Xfpks from *Bifidobacterium breve, Lactococcus lactis* and *Rhodotorula graminis* (Bergman et al., 2016; Evans and Ratledge, 1984; Petrareanu et al., 2014), as well as a phosphoketolase from *Clostridium acetobuytlicum* which favours xylulose-5-phosphate as a substrate over fructose-6-phosphate (Fig 3(C)). Of the four phosphoketolases, only the Xfpk from *B. brevis* (*Bbxfpk*) increased the production of the measured aromatic metabolites significantly, with 2-PE production increasing over 2mM (KmASR.070). However, in all strains, this was accompanied by increased acetate production (Fig. S2 (A)) as well as slower growth compared to KmASR.004. In contrast, overexpressing *KmTKL1* and *KmTAL1* together (KmASR.056, Fig. 3(D)) did not significantly increase production of any measured metabolite except a slight increase in shikimate balanced overall by a decrease in tyrosine.

Finally, we decided to examine which combinations of enzymes best increased both precursors in KmASR.004 by coexpressing *KmENO1* or PEPSes alongside either *Bbxfpk* or the Km*TKL1/TAL1* combination. Relative to our previous attempts to increase phenylalanine production one precursor at a time, only a combination of BbXfpk and KmEno1 further incrementally improved production (KmASR.074, Fig. 3(D)), but not significantly higher than KmASR.070 or KmASR.109. This suggested that the combined effects of enzymes used to increase PEP or E4P supply to the shikimate pathway were not purely additive. Furthermore, neither the PEPSes nor *KmENO1,* when coexpressed with *KmTKL1* and *KmTAL1* in KmASR.004, significantly increased production (KmASR.057-KmASR.059, Fig. 3(D)), nor did coexpressing PEPSes alongside *Bbxfpk* increase production over KmASR.070 (KmASR.070 vs KmASR.071, Fig. 3(D)).

### 3.3. Integrating precursor supply with the phenylalanine biosynthetic pathway

Our engineering of KmASR.004 identified strategies that could increase PEP and E4P supply to the shikimate pathway. We now wanted to test how these strategies would perform in a strain optimised for AAA production, which should exhibit downstream pull through the shikimate pathway. Unexpectedly, the two modifications that had been identified as the best performing in KmASR.004, overexpression of *Bbxfpk* (KmASR.082) by itself or with *KmENO1* (KmASR.072), performed poorly and led to a reduction in titres of 2-PE (and tyrosol and shikimate) when introduced into KmASR.046 (Fig. 4(A)). One possible explanation was that the acetyl phosphate produced by Xfpk is further converted to acetate, lowering the cytosolic pH and increasing metabolic burden (Bergman et al., 2019). Indeed, this appears to be the case because the overexpression of a phosphotransacetylase (Pta, EC 2.3.1.8) from *Salmonella enterica* (*Septa*) (Brinsmade and Escalante-Semerena, 2004) along with *Bbxfpk* alleviated the problem and generated strain KmASR.149 that produced over 4mM 2-PE. We also tested the effects of overexpression of enzymes of the non-oxidative branch of the PPP (Fig. 4(B)). While individual effects of *KmTKL1* and *KmTAL1* were weak, overexpression of both *KmTKL1* and *KmTAL1* (KmASR.062) or the entire non-oxidative branch of the PPP (KmASR.047) yielded ~4.5mM 2-PE. Combining overexpression of *KmTKL1* and *KmTAL1* (KmASR.062) with *KmENO1* (KmASR.063) or bacterial PEPS (KmASR.064, KmASR.065) did not improve the titres of 2-PE (Fig. 4(A)), again illustrating that modifications are not simply additive.

**Figure 4.**
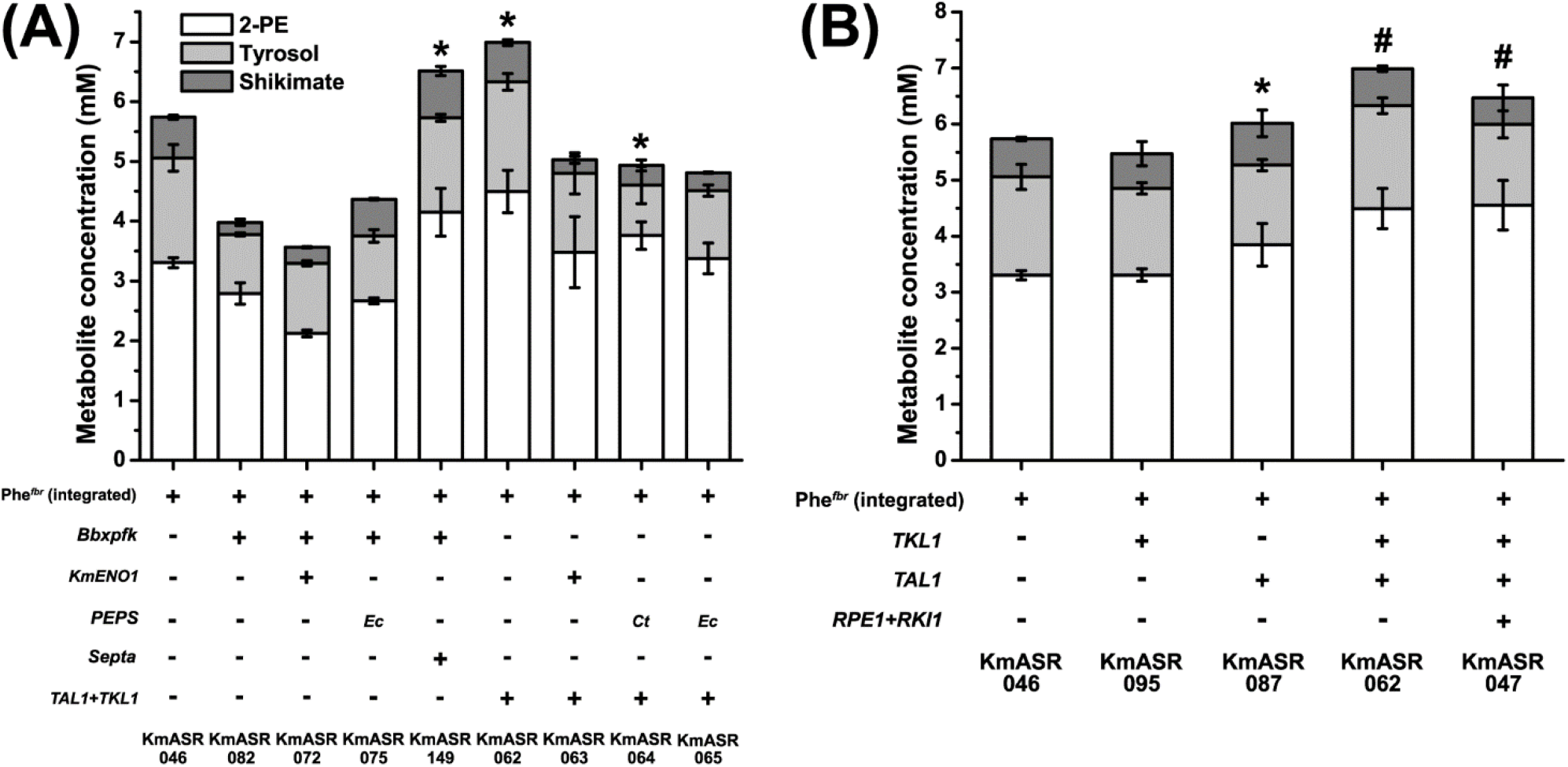
Optimizing phenylalanine production by increasing precursor supply to the phenylalanine pathway. **(A)** Overexpressing heterologous enzymes to increase precursor supply either by themselves or with native enzymes had more variable results, with strategies that worked in KmASR.004 not working in KmASR.046. Ultimately precursor supply via the PPP (KmASR.062) could be matched by using a phosphoketolase (Xfpk) from *B. breve* and a phosphotransacetylase (Pta) from *S. enterica* (KmASR.149). **(B)** Using a partial (KmASR.062) or full (KmASR.047) overexpression of the non-oxidative pentose phosphate pathway (PPP), we were able to increase phenylalanine production in strain KmASR.046 overexpressing a feedback-resistant phenylalanine pathway by 33%. Strains producing significantly higher amounts of 2-PE relative to that produced by KmASR.046 as determined by a paired *t-*test are marked by an asterisk (*, *p*<0.05) or a hash (#, *p*<0.005) All data are plotted as the mean ± s.d. of at least three replicates. Abbreviations for organisms are: *Km: Kluyveromyces marxianus, Ct*: *Chaetomium thermophilum*, *Tt*: *Thielavia terrestris, Bb: Bifidobacter breve, Se: Salmonella enterica*.

Comparing all the derivatives of KmASR.046 tested, KmASR.047, KmASR.062 and KmASR.149 had the most improved production of phenylalanine, all producing up to 4.5mM (550 mg L^−1^) 2-PE. Of these three, we decided to use KmASR.047 as the basis for integrating strategies to simultaneously increase E4P and PEP supply to phenylalanine biosynthesis. While the three mentioned strains produced approximately similar levels of 2-PE, tyrosol and shikimate, KmASR.047 began producing larger amounts of all metabolites within 24h, compared to the other two strains, suggesting that this strain could be better optimized for faster bioprocesses. We now took KmASR.047 as a base strain and considered approaches to improve PEP supply using the knowledge that we had gained. Because we had already found that interactions between pathways were complex, we also retested some of the enzymes that had not performed well previously (Fig. 5(A)). *KmENO1* increased 2-PE production (KmASR.121) whereas overexpressing *EcppsA* or PPDK led to a significant decrease in the production of 2-PE (KmASR.120 and KmASR.142; Fig. 5(A)). Using the PEPSes from *Aspergillus nomius* (*AnPEPS*; KmASR.154) or *Haloferax mediterranei* (*HmPEPS*; KmASR.157) also reduced 2-PE titres but increased shikimate concentrations to ~2mM, compared to 0.46mM in KmASR.047. It may be that these enzymes, catalysing PEP formation reversibly, began to synthesize pyruvate from PEP and deplete levels of the latter, leading to insufficient PEP to carry out the Aro1p-catalysed reaction (Fig. 1).

**Figure 5.**
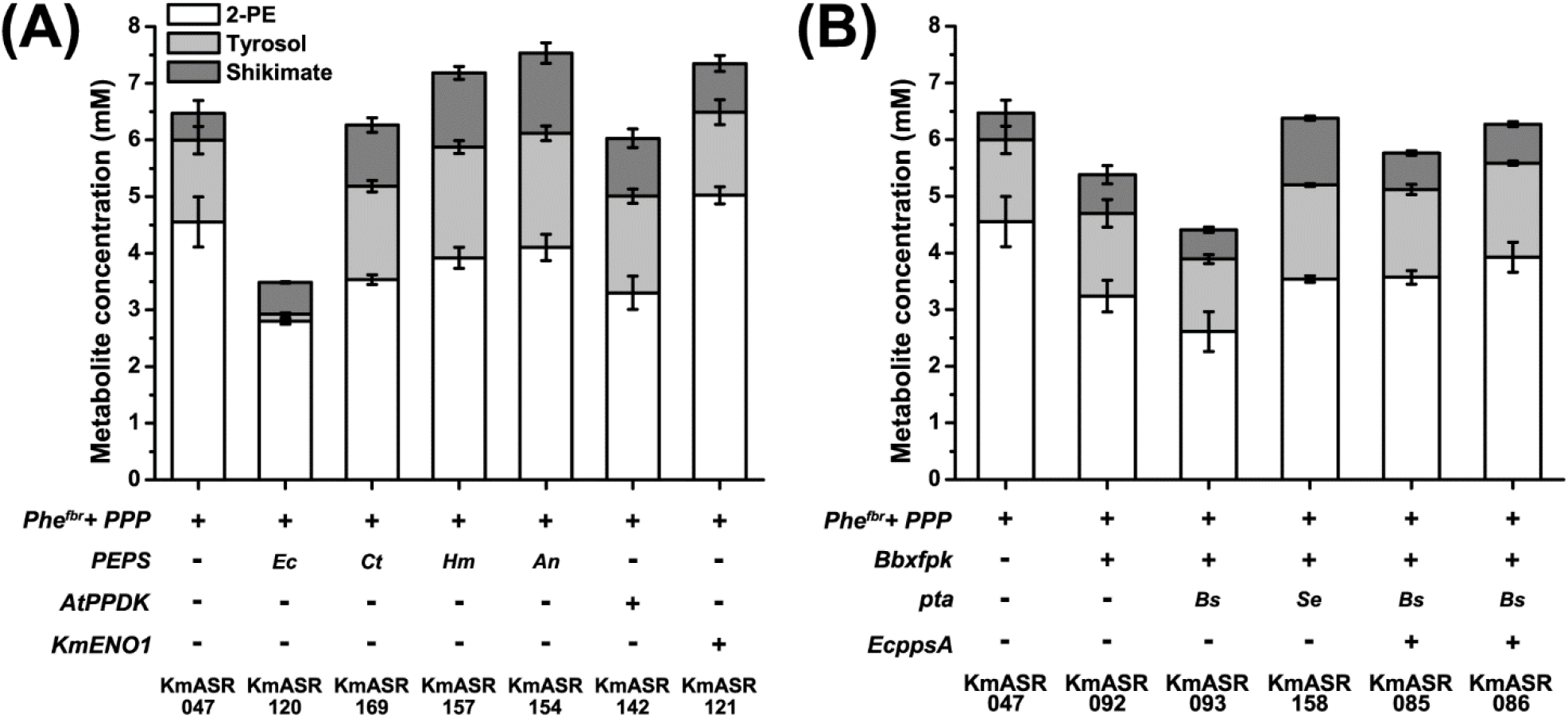
Further integration of precursor supply in a phenylalanine overproducing strain. **(A)** Beginning with strain KmASR.047 with the non-oxidative PPP overexpressed, we attempted to further increase PEP supply using *EcppsA*, the PEP synthase from *E.coli*, a PPDK from *A. thaliana* and the *K. marxianus* enolase, with only the last increasing phenyalanine/-2-PE production. **(B)** Attempts to further increase E4P supply using Xfpk resulted in net decreases in phenylalanine, tyrosine and shikimate production. Expressing a Pta only allowed aromatic compound production to recover if an enzyme was coexpressed to hypothetically simultaneously increase PEP supply. We also found that the Pta from *S. enterica* was more effective in this role. Data are the mean ± s.d. of at least three replicates. Abbreviations for organisms are: *Km: Kluyveromyces marxianus, At: Arabidopsis thaliana, Bb: Bifidobacterium breve, Ct: Chaetomium thermophilum, An: Aspergillus nomius, Hm: Haloferax mediterranei, Bs: Bacillus subtilis, Se: Salmonella enterica*.

Further attempts to increase E4P supply in KmASR.047 using phosphoketolase were also unsuccessful Fig. 5(B)). Overexpressing *Bbxfpk* in this strain decreased 2-PE production by nearly 30% to 3.24mM (KmASR.092) and, in contrast to the previous finding with KmASR.046, this effect was not alleviated by coexpressing *Septa* (ASR.158). We also tested an additional PTA, *Bspta* from *B. subtilis,* but this also did not recover production (ASR.093). An excess of E4P is known to inhibit bacterial DAHP synthase in the absence of PEP (Parker et al., 2001) and could be relieved by addition of excess PEP. We therefore coexpressed *EcppsA* alongside *Bbxfpk* and the two Ptas creating strains KmASR.085 and KmASR.086. While this did improve production over KmASR.093 and KmASR.0158, increasing production overall and converting more shikimate to 2-PE and tyrosol, 2-PE production was still less than KmASR.047. Taken together, these data suggest that careful balancing of the precursors to the phenylalanine biosynthetic pathway must be taken into consideration when increasing their availability.

### 3.4. Optimization of 2-phenylethanol production by knock-down of tyrosine production and aromatic aminotransferase deletion

The 2-PE produced by the best-performing strains so far only amounts to approximately 70% of the total aromatic metabolites produced and 74% of the Ehrlich metabolites of phenylalanine and tyrosine. The third step of our engineering strategy is to knock down expression of the prephenate dehydrogenase *KmTYR1* (Fig. 1). This enzyme converts prephenate to 4-hydroxphenylpyruvate, committing it to tyrosine synthesis: by reducing its expression as low as possible without creating a tyrosine auxotroph, more prephenate from Aro7p should be available for conversion to phenylpyruvate and phenylalanine. To achieve this, we substituted the native *KmTYR1* promoter (*TYR1pr*) with one of two weaker promoters, *GDH2pr* and *REV1pr,* that we had previously characterized (Rajkumar et al., 2019). Both strains, ASR.047kd and ASR.047.kd2 increased the relative proportion of 2-PE to tyrosol, with the strongest effect seen when *REV1pr* (strain KmASR.047kd2) was used Fig. 6). At this stage, 2-PE amounted to 89% of the Ehrlich metabolites and 83% of the aromatic metabolites measured. The decreased tyrosine production was still sufficient for prototrophy, and neither of the knock-downs created a significant growth defect in KmASR.047. Another strategy reported in the literature to increase 2-PE production is to delete the aromatic aminotransferase encoded by *ARO8* (Romagnoli et al., 2015). Inactivation of the putative *KmARO8* gene using CRISPR/Cas9 caused a 10% increase in all measured metabolites, with 2-PE titres increasing to 5.05mM (KmASR.112). We subsequently combined the two strategies in KmASR.047 to examine if these effects were additive for the production of 2-PE (KmASR.117). In KmASR.117, 2-PE production did not significantly increase with the knockout (6.31mM vs 6.25mM in KmASR.047kd2) but formed nearly 93% of the total Ehrlich metabolites, as opposed to 89% and 82% for the *KmTYR1* knockdown and *KmARO8* knockout alone, respectively (Fig. 6). The same modifications, when made to KmASR.062, resulted in a smaller increase in aromatic compound production resulting in the production of 5.2mM 2-PE (~634 mg L^−1^; Fig. S3).

**Figure 6.**
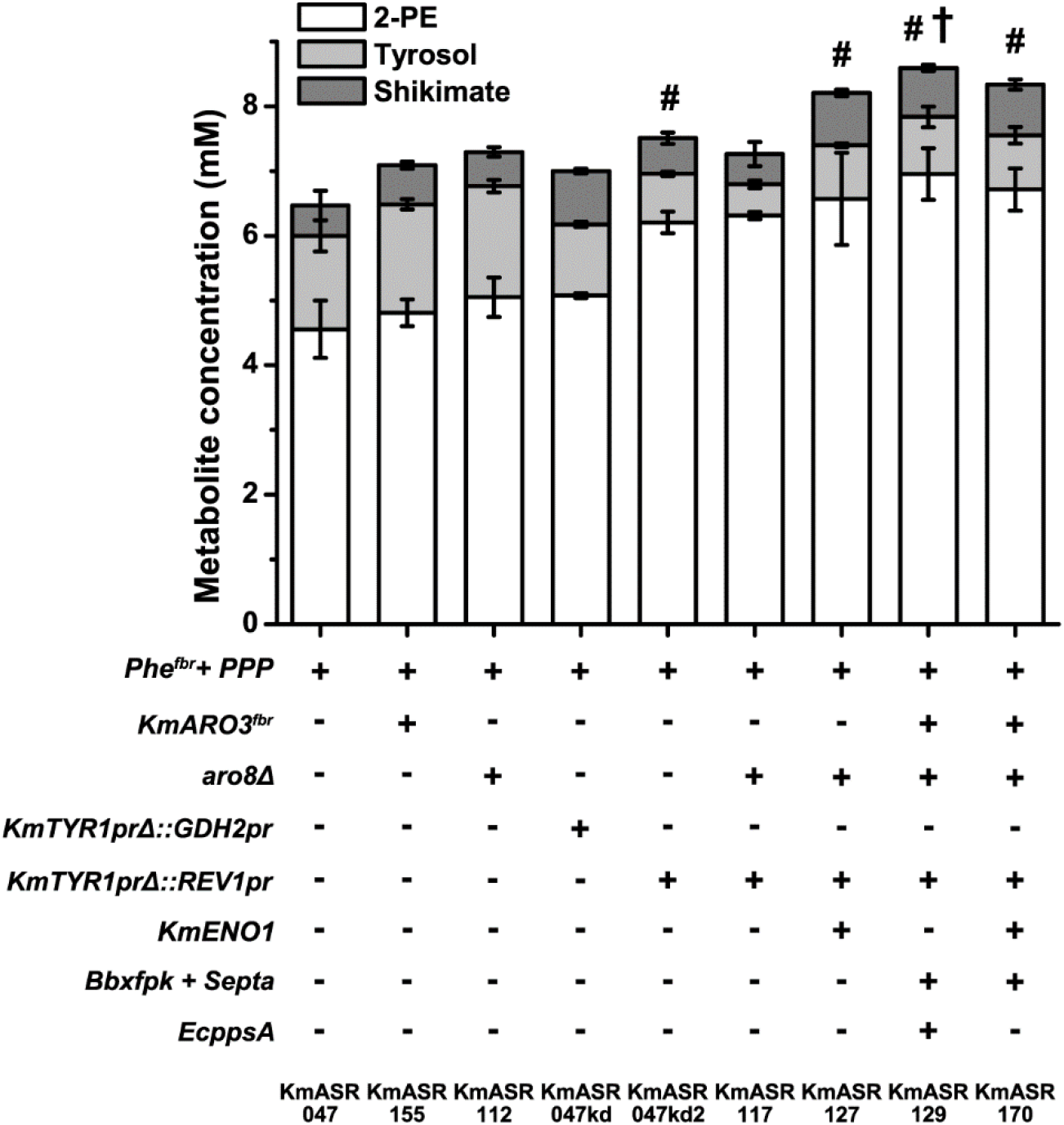
Optimizing phenylalanine production by minimising degradation and competition from tyrosine biosynthesis. KmASR.047’s aromatic amino acid biosynthetic flux was redirected by knocking down *TYR1* expression using a weak promoter (ASR.047kd, ASR.047kd2), and restricting metabolism by knocking out the aromatic aminotransferase *KmARO8* (KmASR.112). The most effective promoter knockdown, the *REV1* promoter, was combined in the latter to produce KmASR.117. With the increased capacity for phenylalanine synthesis in this strain, we increased phenylalanine synthesis more by further supplying precursors using either *KmENO1* or a combination of heterologous enzymes alongside *KmARO*3^*fbr*^ to better utilize increased precursor levels. Data are the mean ± s.d. of at least three replicates. trains producing significantly higher amounts of 2-PE relative to that produced by KmASR.047 as determined by a paired *t-*test are marked by an asterisk (*, *p*<0.05) or a hash (#, *p*<0.005), and those producing significantly more 2-PE relative to KmASR.0117 are marked with a dagger (†, *p*<0.05). Data are the mean ± s.d. of at least three replicates. Abbreviations for organisms are: *Km: Kluyveromyces marxianus, Bb: Bifidobacterium breve, Se: Salmonella enterica*.

### 3.5. Creation of a high 2-phenylethanol – producing strain by a synthesis of tested engineering strategies

The three parts of our engineering strategy so far allowed us to establish feedback-resistant phenylalanine synthesis, optimize E4P supply for phenylalanine synthesis and then redirect metabolic flux within this pathway to create strain KmASR.117. However, lessons learned from other strategies to increase PEP and E4P supply led us to attempt to also increase precursor supply in this strain. Overexpressing *KmENO1* in KmASR.117, led to a further increase in all aromatic compounds produced overall but with 2-PE only increasing by 4% (KmASR.127, Fig. 6). We then decided to take what were now our best strains and see what the effect of modifications for increase of E4P and/or PEP would be. We also included a feedback resistant allele of *KmARO3* since an *ARO3*^*fbr*^ created in *S. cerevisiae* was found to aid aromatic amino acid synthesis when used in tandem with *ARO4*^*fbr*^ (Reifenrath et al., 2018b). Based on the performance of previous strains, we hypothesised that KmAro3^fbr^ would allow faster consumption of the increased E4P and PEP, avoiding their accumulation and diversion to other reactions. The resultant *KmARO3*^*fbr*^ encoding a K222L mutation was overexpressed alongside *Bbxfpk, Septa* and *EcppsA* or *KmENO1* to create strains KmASR.129 and KmASR.170, respectively. Of the two, the best performing strain was KmASR.129, which produced nearly 850 mg L^−1^ (6.95 mM) 2-PE in 48h, the highest titre so far, with a yield of 0.062 mol Phe or 2-PE/mol of glucose. Together, the final yield of Ehrlich alcohols of phenylalanine, tyrosine and shikimic acid from KmASR.129 was 0.077 mol/mol of glucose.

## 4. Discussion

*K. marxianus* is a promising next-generation yeast cell factory on account of its unique physiology, and has begun to be used for such applications. The same physiology and differences in metabolism to *S. cerevisiae* means that metabolic engineering approaches for this yeast require novel, rational approaches to best redirect its metabolites as precursors for valuable chemical compounds. We implemented a three-step rational strategy to optimise a native biosynthetic pathway for overproduction and demonstrated its use for production of a commercially-relevant product de novo from synthetic growth medium. By (i) establishing phenylalanine overexpression via a feedback-resistant biosynthetic pathway, (ii) increasing precursor supply to the pathway, and (iii) knocking down competing pathways, reducing degradation and finally re-balancing precursor supply, our best-producing strains could synthesise 6.95mM, or nearly 850 mg L^−1^, of 2-phenylethanol (2-PE) from glucose in 48h flask cultivation. While we explicitly have engineered *K. marxianus* for the production of 2-PE, relatively few modifications would be required to convert this strain to a phenylalanine production strain. The main modification would involve the inactivation of *KmARO10* and *KmPDC5* to ensure the conversion of phenylpyruvate (PPY) to phenylalanine as opposed to 2-PE. This strain would be a platform for production on phenylalanine-derived products.

Keeping *K. marxianus’* unique physiology in mind, we overexpressed heterologous enzymes which would make use of their high TCA cycle and PPP fluxes to increase precursor supply to the shikimate pathway (Blank et al., 2005; Jung et al., 2016; Sakihama et al., 2019). By screening several PEP synthases, we were able to identify at least two, AnPEPS (in strains KmASR.108 and KmASR.154), and EcPEPS (in KmASR.129) that can effectively convert pyruvate to PEP for the shikimate pathway in strains overexpressing a feedback resistant (fbr) phenylalanine biosynthetic pathway. Although a previous study suggested that they could not function in this manner in *K. marxianus* (Kim et al., 2014), we found that the choice of enzyme and strain engineering played a role in the extent to which PEP synthase overexpression improved AAA or shikimate production. As some PEPSes are exclusively used by their host organisms for either gluconeogenesis or PEP synthesis (Chen et al., 2019; Zhang and Anderson, 2013), a further investigation of more PEPSes with similar functions and kinetics may help us identify the best way to integrate this class of enzyme in yeast cell factories. In our efforts to increase E4P production, we overexpressed phosphoketolases that tap into *K. marxianus’* high fructose-6-phosphate levels and PPP flux (Bergman et al., 2019; Jung et al., 2016) to further boost available E4P. This strategy has recently been employed in other yeasts to increase the production of phenylalanine and tyrosine-derived metabolites with mixed results (Gu et al., 2020; Guo et al., 2020; Hassing et al., 2019; Liu et al., 2019). While it was successful in *K. marxianus*, we found that an Xpfk’s ability to increase the production of aromatic compounds is tempered by the extent of metabolic engineering on the AAA pathway. In contrast to previous studies, we found that coexpressing a Pta with *Bbxfpk* did improve production relative to when just *Bbxpfk* was expressed but that the choice of Pta mattered. (KmASR.149, Fig, 4A; KmASR.158, Fig. 5(B)).

Our final strain modifications highlighted the importance of balancing precursor supply and ensuring that precursors are efficiently utilised. This was especially apparent when optimal strategies to increase E4P and PEP varied across different scales of feedback-resistant 2-PE biosynthesis (Fig. 3, Fig. 5). A combinatorial fine-tuning of gene expression could provide more robust and scalable strategies to increase precursor supply. Other modifications, such as the overexpression of *ARO*3^*fbr*^, only had an effect on 2-PE production when precursor supply was sufficiently increased for the enzyme to optimally function. Finally, as our best performing strains still leaked some shikimic acid from the pathway, overexpression of a bacterial shikimate kinase in addition to *KmARO1* could be considered to ensure all the shikimate synthesised is used downstream and not secreted (Hassing et al., 2019; Rodriguez et al., 2015).

One significant difference to *S. cerevisiae* engineering approaches we found was that whereas in *S. cerevisiae* overexpressing transketolase alone is sufficient to increase E4P levels, in *K. marxianus*, overexpression of both transketolase and transaldolase is required. While besides having a naturally high flux through the PPP, *K. marxianus* has also been reported to have high intracellular levels of fructose-6-phosphate (Jung et al., 2016), one of the substrates for transketolase but for the reverse reaction of interest; these high levels may inhibit the production of E4P by the overexpressed *KmTKL1* and drive equilibrium in the opposite direction. By coexpressing it with *KmTAL1*, the two enzymes may be acting in tandem through their reactions, to ensure E4P production and avoiding the drawbacks of a single enzyme running in the reverse direction. Beyond this, overexpressing the rest of the pathway did not significantly improve phenylalanine production but provided a number of advantages such as a faster onset of production and more capacity for production once competing pathways were eliminated (Fig. S3).

The focus of our study was on developing a rational synthetic biology framework for building a *K. marxianus* chassis to overproduce aromatics. The construction of a cell factory engineered for *de novo* synthesis of 2-PE demonstrated the success of this approach. However, derivatives of a different *K. marxianus* strain (DMKU3-1042) have been reported to overproduce to 1.3 g L^−1^ (10.7mM) 2-PE through a combination of artificial evolution and engineering (Kim et al., 2014), with the host strain natively producing trace amounts of 2-PE.The same study also reported that overexpression of *ARO10* and *ADH2* from *S. cerevisiae* led to a significant increase in 2-PE production. In contrast, in our strain, overexpressing the same genes in wild-type, KmASR.047 or KmASR.117 only had marginal effects (Fig. S4). Based on these data and other indicators, it is likely that underlying differences in AAA metabolism between the two strains affects the effectiveness of particular engineering approaches. Future research into the comparative metabolomics of *K. marxianus* strains will help identify the most metabolically suited strains for specific applications.

## 5. Conclusion

In summary, we implemented a comprehensive synthetic biology strategy to build a *K. marxianus* strain for production of molecules derived from the aromatic amino acid phenylalanine. The potential of this chassis was demonstrated by constructing a cell factory that, starting from zero, produced almost 7mM 2-PE from minimal mineral medium and glucose. Relatively few further modifications would be required to redirect metabolism for alternative products, or indeed to use tyrosine or tryptophan as the starting point. It would also be very interesting to apply our synthetic approach to the use of alternative carbon sources, in particular second generation feedstocks. Our study combined native and heterologous enzymes to increase the supplies of both phosphoenolpyruvate and erythrose-4-phosphate, and while not exhaustive, our findings reveal that balancing the precursors to shikimate and phenylalanine biosynthesis not only affects the amount but also the profile of aromatic metabolites produced. In two key aspects, the importance of synthetic biology approaches were very evident. First, it was necessary to try multiple versions of the same enzyme to identify the most suitable one for a particular reaction; and second, the effectiveness of any given modification was influenced in non-predictable ways by other modifications. Thus, a combinatorial, parallel approach is required to build high-performing strains. Although such strategies have been deployed with *S. cerevisiae* for some time, it was considered that the development of other yeasts was not sufficiently advanced to allow the same approach. With this study, we demonstrate that *K. marxianus* has reached the stage where synthetic biology is feasible and the steps that we followed may serve as a roadmap for other non-traditional, or non-conventional, yeast cell factories.

## Supporting information

Synthetic gene sequences.

Data for all the figures in the text.

## Contributions

ASR and JPM conceived the study and JPM supervised the research. ASR created the expression vectors, engineered the strains, and performed cultures to characterize the strain. ASR and JPM analysed the data, and wrote and revised the manuscript. All authors read and approved the final manuscript.

## Acknowledgements

We thank Javier Varela, Joel Akinola, and Julia Bauer for help with plasmid construction, Else-Jasmijn Hassing for help with the HPLC protocol and Daniel Walsh for technical assistance with HPLC analysis. We also wish to thank Javier Varela, Else-Jasmijn Hassing, Jean-Marc Daran and other colleagues in the CHASSY project for advice and discussions.

