## Supplementary material for "Rational engineering of *Kluyveromyces marxianus* to create a chassis for the production of aromatic products": Synthetic gene sequences.

**Table S5.** Sequences of codon-optimized heterologous genes. The protein sequences were codon-optimized for *S. cerevisiae* using IDT’s online codon optimization tool at <https://eu.idtdna.com/CodonOpt>. Codons creating BsaI or BsmBI sites were altered manually where necessary.

>EcppsA/EcPEPS/A27

ATGTCAAATAACGGCTCATCACCTTTGGTGCTTTGGTACAACCAACTGGGAATGAACGACGTAGATCGTGTTGGGGGGAAGAATGCAAGTCTGGGAGAGATGATCACAAATCTATCAGGGATGGGCGTATCTGTTCCTAATGGATTCGCGACAACCGCGGACGCTTTTAATCAGTTCCTGGACCAATCTGGAGTGAATCAGCGTATTTACGAGCTGTTAGACAAAACGGACATCGATGACGTCACCCAGTTAGCAAAAGCAGGGGCCCAAATTAGGCAGTGGATTATTGACACTCCATTCCAACCCGAACTGGAAAACGCAATCAGGGAAGCCTACGCGCAACTAAGCGCCGACGATGAGAACGCGAGCTTCGCCGTTAGAAGCAGTGCTACAGCAGAGGATATGCCCGATGCCAGCTTCGCGGGCCAACAAGAGACATTCCTAAACGTCCAAGGGTTTGACGCGGTACTTGTCGCTGTCAAACATGTCTTTGCCTCTTTGTTCAATGACAGGGCGATCTCTTACAGGGTCCACCAGGGTTACGACCACAGGGGCGTAGCTCTGTCCGCAGGGGTGCAGAGGATGGTAAGATCTGACCTTGCAAGCTCAGGAGTGATGTTTTCTATTGATACAGAAAGTGGGTTCGATCAGGTGGTATTTATCACTTCTGCCTGGGGCTTAGGTGAGATGGTTGTTCAGGGGGCGGTAAACCCCGACGAATTTTATGTACATAAACCGACTCTTGCGGCTAACAGACCCGCCATCGTTCGTAGGACCATGGGTAGCAAAAAAATTAGAATGGTGTATGCTCCGACGCAAGAACACGGCAAGCAGGTAAAGATTGAGGATGTCCCACAGGAGCAAAGAGACATCTTCTCTTTAACGAATGAGGAGGTTCAGGAACTAGCCAAGCAAGCTGTGCAAATTGAAAAGCACTACGGCCGTCCTATGGATATTGAATGGGCAAAGGACGGTCACACGGGTAAGTTGTTCATTGTGCAGGCCCGTCCTGAAACAGTTCGTTCCAGAGGGCAAGTTATGGAAAGGTACACGCTTCATAGCCAAGGTAAGATTATAGCAGAAGGGAGGGCGATTGGCCATAGGATTGGTGCTGGGCCAGTTAAAGTGATTCACGATATTTCTGAAATGAATAGAATTGAGCCAGGTGATGTTTTAGTTACCGACATGACTGATCCCGATTGGGAGCCTATAATGAAGAAGGCATCCGCCATCGTGACCAATAGGGGCGGAAGGACCTGCCATGCGGCTATAATCGCGCGTGAGTTAGGTATCCCGGCTGTTGTTGGATGTGGTGACGCAACGGAAAGGATGAAGGATGGCGAGAATGTTACCGTATCCTGCGCAGAAGGAGACACAGGATATGTTTATGCAGAGCTGCTTGAATTTAGCGTAAAAAGTTCATCTGTGGAAACCATGCCCGATCTGCCTTTGAAGGTTATGATGAACGTTGGTAATCCCGATAGAGCGTTCGATTTCGCCTGTTTACCCAATGAGGGAGTTGGTCTTGCAAGGCTTGAGTTCATTATTAACAGAATGATCGGAGTCCACCCTCGTGCTCTATTAGAATTTGATGATCAGGAACCACAGTTACAAAACGAAATTAGAGAGATGATGAAAGGGTTCGATAGTCCAAGGGAATTTTATGTGGGCAGATTAACAGAGGGCATCGCAACTTTAGGGGCAGCCTTTTACCCCAAGAGGGTAATTGTGAGACTGTCCGATTTCAAATCCAATGAGTACGCCAATTTAGTTGGTGGCGAAAGATATGAACCAGATGAAGAAAATCCTATGCTGGGATTTCGTGGCGCTGGTAGGTATGTGTCTGATTCCTTTAGAGACTGCTTTGCATTAGAATGCGAGGCCGTAAAACGTGTAAGAAACGATATGGGATTAACCAACGTCGAAATTATGATTCCGTTCGTGCGTACGGTAGATCAAGCCAAAGCTGTTGTGGAGGAACTAGCAAGACAAGGATTAAAAAGGGGGGAAAATGGACTTAAAATTATAATGATGTGTGAAATTCCATCCAACGCCCTTCTTGCCGAACAGTTCCTAGAGTACTTTGACGGGTTCAGTATCGGCTCAAACGACATGACACAACTTGCACTGGGGCTGGACAGGGATTCAGGGGTCGTAAGTGAGCTTTTTGACGAAAGGAACGATGCTGTTAAGGCACTACTATCCATGGCTATCAGAGCTGCAAAAAAACAGGGTAAGTACGTGGGAATCTGTGGACAAGGTCCAAGCGATCATGAAGATTTCGCTGCCTGGTTGATGGAAGAAGGTATTGATTCTCTATCCTTAAACCCCGACACGGTCGTGCAGACGTGGTTGTCCTTGGCAGAACTTAAGAAATAG

>CtPEPS/A26

ATGGCGCCTGACCATTCCACCAACAAGCCATTGGTAAGAAACTTTGAGCACCTGACACGTCAGGATGTGGCGTCTGTCGGTGGCAAGAACTCCAGCCTGGGGGAAATGATTGGAGGTCTTGCATCAAAAGGTGTCGCCATTCCGCCGGGATTTGCCACCACTAGCGACGCGTTTTGGCAGTTTATCGACACGAACTGTCTTAGGGACAAGATCGCCCAGCTTATAGAGCAATGGTCATCCGGTCAAGCGACATTAGCGGAAACGGGGCAAGCTGTACGTAGATTGATAGTCGGAGGCAAATGGCCGGAAGACGCAGCGGCGGCGATAAAAACGGCTTATAGACAGTTAAGCGAGAAGGCGGGTATCAAAAACCTTTCCGTAGCCGTCCGTTCAAGTGCTACAGCAGAAGATTTGCCGTCAGCGTCCTTTGCCGGTCAATTAGAGTCATACCTTAACGTAAGTGGTGAAGACCAATTGCTGGATGCATGTCGTCGTTGTTACGCGAGTCTTTTTACGGACAGGGCTATATCCTACAGGCAGACTCGTGGTTTTGATCACTTGTCCATTGCTCTATCCGTTGGGGTGCAACGTATGGTAAGAGCTGGCGATAGGCACGGCGCGTCAGGCGTCATGTTCAGCCTAGACACCGAAAGCGGCTTTGATCAAATAGTCTTAATAAACGCGGCCTGGGGCCTGGGGGAAAATATAGTCCAGGGCGCAGTAGACCCGGACGAGTACCAGGTATTCAAACCTCTGTTGAATAACCGTCCAGACCTAGTTCCGATTGTACAAAAGAAGAGAGGCGAAAAAGAAATAAAAATGATCTATGGTGATGGGGAGACTATTCGTACTAGAAACGTGCCCACCTCCAGGGCCGAAAGGGCCGCATTTGTGTTGAGCGATGAAGACATATTGCAGCTTGCTCGTTGGGCCTGTACTATCGAGAAGCATTACGGGTGTCCGATGGATATGGAGTGGGCCAAGGATGGTACTACAGGCGAATTATTTATCGTCCAAGCCAGACCAGAAACGGTGCATAGTAGAAACACTTCTGCCGTTATTAAGACGTACGACGTAGGCAAAAAAGGCAGGATATTGGCGACGGGCCACAGCATTGGCGATGCTGCTGTAACTGGCCGTGTGTGTATGATAGAAAGTGCAAAAGATATTGATAAGTTCGTCGAGGGGTCTGTCCTAGTGACAGGCAGTACCGATCCAGACTGGGTGCCCATTATGAAAAAGGCGGCCGGTATTGTCACCGACCATGGGGGACGTACTTCACATGCAGCTATCGTCAGTCGTGAATTAGGCTTACCTGCTGTAGTCGGAACAGGTAATGCCACTTACGTGCTGCACACGGGGCAAGATGTGACAGTGAGCTGCGCGGAGGGGGACATCGGTTATGTCTACGAAGGCGTGTCAGAGATTACGACACATACAGTAGATCTAGCAAAAATGCCTACGGTGCAAACAAATGTCATGTTAAACCTGGCCAACCCTGCAGCAGCCTATCGTTGGTGGAGGCTGCCGTCAGACGGTATAGGACTAGCTAGAATGGAGTTTGTTATAAACAATGCTATATGCGTTCATCCTATGGCCCTGGTTAGATACGACCAGTTAAAGGACGAAGACGCTAAGAAGGAGATTGCGAAGCTTACAATCGGGTATACGACCAAGCCCGACTACTTCGTGGATACTCTGGCAAGGGGTCTGGCCTCATTGTGCGCAACGGTCTACCCTAGACCCGCCATTATTCGTATGAGTGACTTTAAAACGAACGAATACGCAGGATTACTTGGTGGCGATGAATTCGAGCCGAGAGAAGAAAACCCAATGCTTGGCTTTAGAGGTGCTTCCAGGTATTATTCACCTCAGTACAAAGAAGGATTCGCTCTTGAGTGCAGAGCCATTAAGAAATTAAGGGAAGTCCTAGGGTTTACTAACGCCATAGTTATGATCCCATTCTGCAGAACTGTCGCGGAGGCGAAGAAGGTCCTAGACGTGATGGGGGAGAACGGGTTGAAACGTGGGGAGAACGGCCTTCATGTATACGTCATGTGCGAGATTCCCTCAAATGTTATACTAGCCGCCGAGTTTTGTAAGTATTTTGACGGGTTCTCAATAGGGAGCAACGATCTAACGCAACTAACATTAGGAGTCGACCGTGACAGTCGTGAGTTAGCAGACCTATTTAGCGAGCAAGATGAAGCGGTGAAGTGGATGATTGCGAGAGTAATCAGCGTCGCTAGAAAAGAAGGACGTAAGATCGGCATCTGTGGACAAGCTCCATCCGACCATCCCGAATTCGCCAAGTTTTTAGTGAGTGCAGGGATTGATAGCATCAGTGTTTCCCCTGATTCCTTTCTTCCAGTGAAACAGAATATTGTAGAAGCAGAGGGCGCATAG

>TtPEPS/A28

ATGAGTGCAAGTAACGAAGATGCCAACCCTCTAGTGTGTGATTTCAAACATTTACTTCGTTCTGATGTAGGTTTGGTAGGAGGTAAGAATAGCAGTTTAGGTGAAATGCTTAGCGCATTAAGTAGTAAAGGAATCGCTGTCCCCCCAGGGTTCGCCACCACTAGCCACGCTTATTGGCATTACGTGGACGCTAACGGGATAAGAGACAAGATCGGTCCGTTAATTTCCGAATGGCAAGCGGGTCACGCTTCTCTGGCTGAGACAGGTCAAGCTGTCAGACGTCTATTCCTGAGAGGCAGCTGGCCAGCGGATGCAGCAGAGGCAATCACAACGGCGTACCAGAAACTGAGCGCCAAAACAGGGGTTGAAAATCTTAGGGTGGCTGTGAGGAGTAGCGCCACAGCAGAAGACCTTCCCGATGCATCCTTTGCGGGCCAATTAGAGAGCTACCTGAATATCTGGGGTAAAGATAGGCTTCTGGACGCCTGCAGACGTTGTTATGCATCCTTGTTTACTGATCGTGCTATTAGTTATCGTCAAGCTAAAGGTTTTGACCATATGTCAATAGCGCTGTCAGTTGGGGTGCAGAGAATGGTTAGGAGCGACGCTTGGGGCAGTGGTGTTATGTTCAGCATCGACACAGAGTCAGGCTTTGACAAAATTGTGTTGATAAATGCCGCTTGGGGTTTAGGTGAAAATATTGTACAAGGTACAGTGAACCCTGATGAATACCAGGTGTTTAAGCCGCTATTGGACGACCCGTCCCTGGTGCCCATAATCCAGAAGAAGCGTGGCGAAAAGGCCATGAAGATGATTCTAGGCCGTTCTCACCACCACCATCCTCATGCACCCACTAGGAACGTTCCCACCTCCAAGGCGGAGAGGGCAGCTTTTGTGCTAGCTAACGATGAAATTCTGCAACTGGCTAGGTGGGCTTGCGCAATAGAAAAGCATTACGGATGTGCCATGGATATGGAGTGGGCCAAAGATGGGACCACCGGTGAACTTTTTATTGTACAAGCTCGTCCAGAAACCGTTCACTCAAGGCAAGACAGCGCCGTATTTAAAACCTATACGGTAAAGAACAAGAGTAGGGTACTAGCGACAGGCTTGTCCATAGGCGATGCCGCCGTATCCGGTCAGTTGTGCTTGATAGAGGACGTTAAGGACATTGGTAAATTTGTAGATGGTTCCATCTTGGTAACGGTTAGCACGGATCCTGATTGGGTACCGATTATGAAGAGGGCAGCGGCCATTATAACCGATCACGGCGGCCGTACATCCCACGCGGCAATTGTGTCCAGGGAGCTGGGTGTTCCAGCCGTGGTGGGAACTGGTGATGCAACATATGTCCTTCATACTGGACAAGATGTAACTGTTTCTTGCGCAGAAGGTGATGTTGGTTTTGTATACGCCGGAATTTCTGACATAGTCGCCACCACTATTGATCTTAAAGGCCTGCCGGCGGTCCGTACAAACGTTATGCTGAACCTAGCGAATCCGAGTGCGGCGTACCGTTGGTGGAGACTTCCCGCCGACGGAATTGGGCTTGCCCGTATGGAGTTTGTTGTTACGAATGCCATTAGAGTTCATCCGATGGCTCTGGTCAGATTTGACAGGTTGCAGGACCAAGCCGCTAAAGCCGAAATTGCTCGTCTTACTGCCGGGTATGAGCATAAGCCAGATTACTTCGTAGACAAACTTGCCCACGGGTTAGCCGCTCTTTGCGCTACGGTCTACCCCAAACCAGCTATCATTAGAATGTCCGATTTTAAAACAAATGAGTACGCGGGTCTAGTAGGTGGAGCGGAGTTTGAGCCAGCCGAGGAGAACCCTATGTTAGGGTTCAGAGGGGCGTCACGTTACTATAGTCCTCGTTATGCAGAGGGCTTCGGTCTAGAGTGCCGTGCAATAAAGTTGTTGAGGGAGGAGATGGGCTTCGCGAATGCCGTGGTTATGATTCCATTCTGCAGAACTGTCGGCGAGGCGGAAAAAGTCTTGCGTGTCATGGAGCAAAACGGGTTAAAAAGGGGCGAAAAGGGTTTGAAAGTTTACGTAATGTGTGAGATCCCTTCTAATGTAATTTTGGCCGAGAGGTTCACAGAGCACTTTGATGGATTTTCTATCGGGAGTAACGATTTGACACAGCTTACGTTAGGAGTCGACAGAGACAGCGGCGAATTAGCCGAATTATTCGATGAACAGGATGAGGCGGTTAAATGGATGATAGCCAAAGTCATCGAAGTTGCGAGGAAGAAAGGGTGCAAGATAGGTATTTGTGGACAAGCACCCAGTGACCACCCGGAGTTCGCGAAGTTCTTGGTTCAAGCCGGCATTGATAGTATCTCTGTGAGTCCCGATTCCTTCCTGGCGGTAAAGAGGCACGTTGTAGACTCTGAAAAGGCTTAG

>AtPPDK/A29

ATGATGCAAAGGGTATTCACGTTTGGTAAGGGCAGGAGTGAGGGGAATAAAGGAATGAAGTCCTTATTGGGAGGCAAGGGAGCGAACCTGGCCGAGATGGCATCAATAGGTCTGAGTGTGCCACCTGGGCTGACCATTTCTACGGAAGCTTGTCAGCAATATCAGATAGCTGGGAAGAAGCTACCAGAAGGATTGTGGGAAGAAATACTAGAAGGACTATCTTTCATTGAGAGAGACATCGGGGCCTCACTAGCGGATCCCTCTAAACCGTTATTATTATCTGTCCGTTCTGGCGCGGCGATCAGTATGCCGGGGATGATGGATACAGTGCTGAATTTGGGTCTTAATGACCAGGTAGTTGTTGGTTTGGCCGCAAAGTCTGGAGAGAGGTTTGCGTATGACAGCTTTAGAAGGTTCTTGGATATGTTCGGAGATGTTGTTATGGGAATTCCTCATGCTAAGTTCGAGGAGAAGCTAGAGAGGATGAAAGAGCGTAAGGGGGTTAAAAACGACACTGATTTAAGCGCCGCTGACCTTAAAGAGCTGGTTGAGCAATACAAGTCCGTTTATCTTGAGGCTAAGGGCCAGGAATTTCCTTCTGATCCGAAAAAGCAGTTGGAGCTTGCGATAGAGGCGGTGTTTGATAGTTGGGACTCACCAAGAGCAAATAAATACAGATCAATCAACCAAATTACTGGACTAAAGGGCACGGCTGTGAATATTCAATGCATGGTATTCGGCAACATGGGTGATACGAGTGGAACTGGGGTGTTGTTCACACGTAACCCGTCCACAGGAGAAAAGAAACTATACGGAGAGTTTCTGGTGAACGCCCAAGGTGAGGATGTAGTAGCAGGAATACGTACACCAGAGGACTTAGACACCATGAAAAGATTTATGCCTGAAGCATACGCTGAGTTGGTAGAAAATTGTAATATCCTGGAGAGGCATTATAAAGACATGATGGATATTGAGTTTACTGTTCAGGAGGAGAGGTTGTGGATGCTACAATGCAGGGCAGGAAAGAGGACTGGGAAGGGGGCAGTGAAAATCGCGGTGGATATGGTGGGTGAAGGTCTGGTGGAGAAAAGTAGTGCCATTAAAATGGTCGAACCTCAGCACCTTGACCAGCTGTTACACCCTCAGTTTCATGACCCCAGTGGCTACCGTGAAAAGGTAGTAGCCAAAGGGCTGCCCGCTTCCCCCGGTGCAGCGGTCGGTCAAGTGGTATTTACTGCGGAAGAGGCTGAGGCGTGGCACTCTCAGGGCAAGACGGTAATCTTGGTTAGAACGGAAACATCTCCTGACGATGTGGGAGGAATGCATGCGGCTGAGGGAATCTTAACTGCCCGTGGTGGTATGACTAGCCACGCCGCAGTCGTAGCTAGAGGATGGGGTAAGTGCTGCATCGCCGGCTGTTCAGAGATACGTGTAGACGAGAACCATAAGGTACTACTTATCGGGGACTTAACGATCAATGAAGGGGAATGGATATCAATGAACGGTTCCACAGGGGAAGTGATCTTGGGCAAACAAGCGCTTGCCCCTCCTGCACTGTCTCCAGACCTTGAAACATTTATGTCTTGGGCTGATGCAATTAGAAGACTTAAGGTCATGGCAAACGCGGACACACCAGAGGACGCCATCGCAGCGCGTAAAAATGGAGCCCAAGGGATAGGGCTGTGCAGGACGGAGCACATGTTTTTCGGGGCAGACCGTATAAAAGCAGTGAGGAAAATGATTATGGCAGTGACAACTGAGCAACGTAAAGCAAGCCTGGATATATTGTTACCATACCAACGTTCCGATTTCGAAGGTATTTTTAGAGCGATGGACGGCCTACCCGTCACAATAAGGTTATTGGACCCTCCTCTACACGAGTTCCTGCCGGAAGGGGACTTAGATAACATAGTACACGAACTGGCGGAGGAAACTGGAGTCAAAGAAGATGAGGTGCTATCAAGAATTGAAAAGTTATCCGAAGTAAATCCAATGCTGGGTTTTAGGGGTTGTCGTCTAGGAATATCATACCCTGAGTTGACTGAGATGCAGGCTAGAGCAATTTTTGAAGCAGCCGCTTCCATGCAAGACCAGGGTGTAACAGTGATCCCAGAAATCATGGTTCCGCTTGTCGGCACTCCTCAAGAGTTAGGACACCAGGTAGACGTTATTCGTAAAGTAGCTAAGAAAGTGTTCGCAGAAAAGGGTCACACTGTATCATATAAAGTCGGAACGATGATAGAGATTCCTAGAGCTGCGCTAATCGCAGACGAGATAGCCAAAGAAGCCGAGTTTTTTTCTTTCGGCACTAACGATCTTACCCAGATGACATTCGGATACAGTCGTGACGATGTTGGCAAGTTTTTGCCAATATATCTGGCTAAAGGTATACTTCAACACGACCCCTTTGAAGTCCTGGATCAACAGGGAGTTGGGCAACTAATCAAAATGGCAACCGAAAAAGGTCGTGCAGCCAGGCCTAGTCTAAAGGTTGGGATATGCGGGGAACACGGGGGGGATCCTTCCTCAGTGGGTTTTTTTGCTGAAGCCGGGTTGGATTATGTTAGTTGTAGCCCCTTCCGTGTACCGATTGCTAGACTAGCTGCCGCCCAGGTCGTAGTCGCGTAG

>AnPEPS/A37

ATGAATGATGCATCTGCTCATGCAAAATTCGTCAGGAATTTCGAGCAGCTTAAAAGAGAGGATGGTCCCTTAGTCGGCGGAAAGAACAGCAGTCTAGGCGAGATGATAACTGCATTAGGGGGGAAAGGTATTGCCGTACCACCAGGCTTTGCTACAACGTCTAGTACTTACTGGCATTATCTTGATGTGAACGACATCCGTAAGAAAATTACCAAATTGATTGAGAACTGGCAGCTAAAGAAAACTACTCTAGCGGAAACCGGCCACGCAGTTAGGGCACTTTTTCTTCATGGAAATTGGCCGACAGACGCAGAAGCCGCTATAAGAGCGGCATATAGACAACTGTGCGCTAAAGCGGGCGTTGATGACTTGTCTGTCGCTGTCAGATCTTCCGCAACAGCGGAGGATCTACCCGATGCTTCTTTTGCAGGTCAACAGGAAAGCTACCTTAACATTAGTGGAGAGGAGGCCCTATTACATGCCTGCAGGCGTTGTTACGCCTCCCTTTTCACTGATAGAGCCATATCTTACCGTCAAGCTAAAGGTTTCGATCACATGAGTGTTGCTTTGTCCATCGGAGTGCAGAGAATGGTCCGTTCTGATGTAGGTGGGAGTGGTGTGATGTTCTCCATCGACACAGAAACGGGATTTGATAAAGTTATTCTAATTAACGCAGCTTGGGGACTGGGAGAAAATGTTGTGCAGGGCACTGTAAACCCTGACGAGTATCAGGTTTTCAAACCTCTGTTAGCTGATCAAAGTTTGGTACCTATAATAGAGAAAAAAAGGGGAGAAAAGGGCATGAAAATGGTCCTAGGCGGGACAGAGACACCCACCAGGAACCTTCCTACATCAAAATCAGAACAGGCTTCTTTCGTCTTGAATGATGGTGAGATCCTACAACTGGGTAGGTGGGCCTGCACAATTGAAACACACTACGGGTGCGCAATGGACATGGAGTGGGCTAAGGATGGTATAACTGGCGAGTTATTTATAGTGCAAGCACGTCCCGAGACAGTACATAGCAGATCACACGCAGCGATATTTAAGACTTATAAGGTGGGAAAAAAAGGGAGGCTATTAACTACCGGGCTGTCTGTGGGCGATGCGGCAGTTTCTGGAAGGGTCTGCTTAATAGAAACAGCCAGGGACATTGATAAGTTTATCGATGGCAGTATACTGGTCACAGAAACAACAGACCCTGATTGGGTTCCGATCATGAAAAGAGCGGTCGCGATAATAACCGACCACGGTGGCCGTACAAGCCACGCTGCTATTATCTCCAGAGAATTGGGACTGCCGGCAATAGTCGGTGCGGGAAACGCAACGTACATTCTGCATACGGGTCAGGACGTTACGGTGTCATGCGCTGAAGGTGACAACGGTTTTGTCTACGAGGGGAGTTCTGAAATTACAACTGAATCTGTTGATTTGACGGAATTCCCAGAAACAAGAACGAAAATTATGCTGAATTTAGCTAACCCCGCCGCGGCATTTAGATGGTGGCGTTTGCCTGCAGACGGGATCGGTCTAGCTAGAATGGAGTTCGTAGTGACTAACGCCATCCAAGTTCACCCAATGGCACTAGTACATTTCGATCAGTTAAGAGATGCAAAAGCCAAAGAGGATATAAGGAGATTGACAACGGGGTACGACAATAAACCCGAATACTTTGTTGATAAACTTTCACATGGGTTCGCTTCACTATGCGCTGCAGTGTATCCGAAGCCTGCCATCATCCGTATGTCAGATTTCAAAACTAACGAATATGCGAGATTAGTGGGAGGAAGAGAATTCGAACCGGACGAAGAAAACCCTATGCTGGGCTTCCGTGGGGCCTCAAGGTACTATAGCGCACACTACAAAGAGGGATTCGCTTTGGAATGCCTAGCCATCAAGAGGCTGAGAGAGAGAATGGGTTTTAGAAATGCGATTGTGATGATTCCTTTCTGCAGAACGGTGGGCGAGGCTAAGAAAGTACTAGATGCGATGGCGGAGAACGGTCTAAGGCGTGGAGAAAATGGCCTACAAGTGTATGTTATGTGCGAGATTCCCTCAAATGTTATCTTAGCGGCGGACTTTATGGAATATTTTGACGGTTTCTCAATAGGTAGCAACGATCTTACACAATTAACTCTAGGTGTGGACAGGGACTCCGGCGAGCTGGCAGATCTATTCGACGAACAAGATAAAGCAGTTAAATGGATGTTACAGCAAGTGATACAAGTAGCCAAACAGAAAGGATGTAAAATAGGAATATGTGGACAGGCGCCTAGTAATCACCCGGAATTCGCGCGTTTTCTAGTGCAGTGTAAGATCGACTCTATATCAGTGTCTCCCGATTCATTTTTTGCGGTGAAGAGGCACGTCGTCGCAGGTGAAAATACCTAG

>HmPEPS/A39

ATGGCAGTCGTTTGGTTAGACGACGTTAGAGCCGACGATTTAGATTTAGTCGGCGGGAAAGGTGCATCCTTAGGTGAGCTGACAGGTGCCGGTTTGCCGGTGCCTCCAGGATTTGTCGTAACTGCATCTACATACAGAGCATTCATAGAAGATGCAGGCATAGACGATGAATTGTTTAGCGCGGTTGAAGTTGACCATGAAGATTCAGCCGCTTTGAAGGAAGCACACGAAGCAGCTCATGAACTAATCATGGGAACACCAATTCCCGATGATGTCAGAGAAGAGATTTTGGATGCTTACAGATCCATTGGTGAAGGGGATGCATTTGTCGCTGTTAGATCGAGCGCGACAGCCGAGGACCTTCCCGATGCGTCTTTCGCGGGACAACAAGAAACGTTTTTAAATGTTACCGAAGACGAGCTGTTGGAAAGGGTTAAAGAATGTTGGGCCTCCCTATTTTCGGAGAGGGCCATATACTATAGAAACCGTAAGGGATTTCCCCACGATAAAGTTGATATCGCAGTTGTGGTTCAACAAATGGTAGACGCAGAGAAATCCGGTGTTATGTTCACGAGGCACCCAAGTACTGGAGATATGAAAGTGATAATTGAGGCCGCTTGGGGGTTAGGCGAGGCCGTTGTTTCTGGCACAGTCAGTCCAGATAATTACGTGGTTGGTAGAGAAGGAGGTGAAGTTGAAACTGCTACAATAGCTGATAAAAAAACCATGTGCGTCAGGGACGAAGAAACTGGTGAAACCGTAATGAGAGATGTCCCAAATGATAGGCGTCACGATAGAGTATTGACCGATTCAGAAGTCGATCGTTTGTTAGAATTGGGAGAACTTGTTGAAGATCATTACGAATCTCCTCAAGATGTTGAGTGGGCCGTCTACGATGGTGATGTTTACATGTTACAATCTAGACCAATCACTACAATTAGTGAAGATGGGGGCGGGGAATCTGCAGGTGAATCAAAAAGATCGACGGAAGCGGGCGATGATGTCATTATGCGTGGTCTTGGTGCCTCACCAGGTATAGCAAGCGGTAGAGCGAGAACAGTTACCAAGCTAGATCATCTAGATCAAGTTGCCGAGGGTGATATTATCGTTACCGAGATGACTATGCCCGATATGGTGCCAGCAATGAAACGTGCTGCAGGGATCGTCACAGATGAGGGTGGTATGACTTCCCACGCTGCAATCGTAAGTAGAGAATTGGGCGTTCCTGCAGTTGTGGGCACCGGAGATGCCACTACGACCCTTGATGATGGAGAAATCGTAACAATAGATGGCGATAAAGGTACCATTAGAGAAGGTGAGGAGCCTGAAGAAAAGCAAAGACAACCTGTTGAGGAAGCGAGGCCTAAGACCCCCGTTAAACCTATGACTGCAACTGAAGTTAAGGTTAATGTTAGTATTCCTGAAGCTGCCGAAAGAGCTGCTGCCACAGGAGCCGATGGAGTTGGTCTATTGAGAATAGAACATATGGTTTTAACTCTGGGAAAGACACCCGAAAGGTTTATAGAACAGAACGGCGAAAGAGCGTATGTCGATGAAATAGTAAGCGGTGTTAGGGAGGTTGCTGAGGAATTTTACCCTAGACCTGTGCGTGTGAGAACGTTGGATGCCCCAACGGATGAGTTCAGGCAACTTGAGGGTGGCGAAAACGAACCAAGGGAACATAACCCTATGTTAGGGTACAGAGGTATCAGAAGAGGTTTGGATAACCCAAATAATTTTAGACTTGAGTTACAAGCGTTCAGAAGATTATTCGACATGGGTTACGATAATGTTGAAATCATGTTCCCATTAGTTAATGATGCCGAAGATGTATTAAGAGCAAAAGAACATATGATTGAGGCAGGTATAGACCCGGAAAAGAGGGAATGGGGTGTTATGATAGAAACACCTGCGGCTGCATTGGGAATAGAAGAAATGGCTGATGCTGGTATAGACTTTGCCTCATTTGGAACCAATGATTTAACACAGTACACTCTGGCAGTAGATAGGAATAATGAACACGTCGCTAACTTATATGATGAACTACACCCTGCTGTATTAAAGTTGATCGGGGACACAGTGGATGCATGCAGGGAAGTCGGTGTTAAAACCTCGATTTGTGGCCAGGCCGGAAGCAAGCCAAAGATGGTGCAATTTCTGGTGGACAAGGGTGTATCATCTATTTCTGCCAATATTGATGCAGTTAGGGATGTACAACATGAAGTGAAGCGTGTTGAACAGAGATTAATTTTGGATTCGGTTAGGTAG

>Bbxfpk/B18

ATGACTAACCCTGTAATCGGTACTCCTTGGCAAAAGTTGGATAGACCAGTTTCAGAAGAAGCAATCGAAGGTATGGATAAATATTGGAGAGTTACCAACTATATGTCCATAGGTCAAATCTACTTGAGAAGTAACCCATTGATGAAGGAACCTTTTACTAGAGATGACGTTAAGCATAGATTAGTCGGTCACTGGGGTACTACACCAGGTTTGAACTTCTTGTTGGCCCATATCAACAGATTGATCGCTGATCACCAACAAAACACCGTTTTTATAATGGGTCCAGGTCATGGTGGTCCAGCTGGTACTTCCCAAAGTTATGTTGACGGTACTTACACTGAATACTACCCAAACATAACAAAAGATGAAGCTGGTTTGCAAAAGTTTTTCAGACAATTCTCCTATCCAGGTGGTATCCCTAGTCATTTTGCACCAGAAACCCCTGGTTCAATTCACGAAGGTGGTGAATTGGGTTATGCTTTATCTCATGCTTACGGTGCAGTAATGAATAACCCATCATTGTTTGTTCCTTGTATTATAGGTGACGGTGAAGCCGAAACAGGTCCATTAGCTACCGGTTGGCAATCTAACAAATTGGTCAATCCAAGAACTGATGGTATCGTATTGCCTATCTTGCATTTGAACGGTTACAAGATTGCAAATCCAACAATCTTGGCCAGAATATCTGATGAAGAATTACATGACTTTTTCCGTGGTATGGGTTATCACCCTTACGAATTTGTTGCCGGTTTCGATAATGAAGACCACATGTCTATCCACAGAAGATTCGCTGAATTGTTCGAAACTATCTTCGATGAAATTTGTGACATAAAAGCTGCTGCTCAAACCGATGACATGACTAGACCATTCTACCCTATGTTGATTTTTAGAACTCCAAAGGGTTGGACATGCCCTAAGTTCATCGATGGTAAAAAGACAGAAGGTTCCTGGAGAGCACATCAAGTTCCATTAGCTAGTGCAAGAGATACCGAAGAACACTTTGAAGTCTTGAAAGGTTGGATGGAATCTTACAAGCCTGAAGAATTATTCAATGCAGATGGTTCAATTAAAGATGACGTTACAGCCTTTATGCCAAAGGGTGAATTGAGAATAGGTGCCAATCCTAACGCTAATGGTGGTGTTATCAGAGAAGATTTGAAATTGCCAGAATTAGACCAATATGAAGTAACTGGTGTTAAGGAATACGGTCATGGTTGGGGTCAAGTTGAAGCCCCTAGAGCTTTGGGTGCATATTGTAGAGATATCATTAAAAATAACCCAGACTCCTTTAGAATATTCGGTCCTGATGAAACAGCTAGTAACAGATTGAACGCAACTTATGAAGTAACCGATAAGCAATGGGACAATGGTTACTTGTCTGGTTTAGTTGATGAACACATGGCAGTCACTGGTCAAGTAACAGAACAATTATCAGAACACCAATGCGAAGGTTTCTTGGAAGCATATTTGTTAACAGGTAGACATGGTATTTGGTCTTCATACGAATCTTTTGTACATGTTATCGATTCAATGTTGAACCAACACGCCAAATGGTTAGAAGCTACTGTTAGAGAAATACCTTGGAGAAAGCCTATCTCCAGTGTTAACTTGTTAGTCTCTTCACATGTATGGAGACAAGATCATAATGGTTTTTCTCACCAAGACCCAGGTGTCACATCATTGTTGATTAATAAGACCTTCAATAACGATCACGTTACCAACATCTATTTTGCCACTGACGCTAACATGTTGTTGGCTATCTCTGAAAAGTGCTTCAAGTCAACTAACAAAATCAATGCAATATTCGCCGGTAAACAACCAGCACCTACATGGGTTACCTTGGATGAAGCCAGAGCTGAATTAGAAGCTGGTGCTGCTGAATGGAAATGGGCTTCTAATGCAGAAAATAACGATGAAGTTCAAGTTGTCTTGGCATCCGCCGGTGACGTCCCAACACAAGAATTGATGGCCGCTAGTGATGCTTTGAACAAAATGGGTATTAAGTTTAAAGTAGTTAACGTCGTAGATTTGTTGAAGTTACAATCAAGAGAAAACAACGATGAAGCATTGACTGACGAAGAGTTTACTGAATTGTTTACTGCTGATAAACCAGTATTGTTTGCATATCATTCCTACGCCCAAGATGTTAGAGGTTTGATCTATGATAGACCAAACCATGACAATTTCCACGTTGTCGGTTACAAAGAACAAGGTTCAACCACTACACCTTTTGATATGGTCAGAGTAAATGATATGGACAGATATGCATTGCAAGCAGCCGCTTTGAAGTTAATTGATGCAGACAAATACGCCGATAAGATCGACGAATTAAACGCTTTTAGAAAGAAAGCATTTCAATTCGCAGTCGATAATGGTTATGACATTCCAGAGTTTACTGATTGGGTATACCCTGATGTTAAGGTTGATGAAACACAAATGTTGTCTGCTACTGCTGCTACTGCTGGTGACAATGAATAA

>Rgxfpk/B17

ATGGCAGATCACCAGGACGCTCCGCCCCCGCCCATCACACCCTCTTTGTACGCGGACAAACCGGATGCACCTCTTTCAAGTTTGCCAGTGCAGCTTGATGTCGATAGTTTGGTAAAAAAGCTACCAGAGCAACACTTGGAAGCTATAGCTGGCAACTGGCGTTTGTCTTGCTACTTGGGAACGGCTCAGACCTTTCTGCAAAAGAATGGACGTTTAGCCCGTAAACTGGAGGTCAGCGACATCAAGCCCCGTTTACTGGGGCACTTAGGTACGCAAGGAGGGCTGTCATTAGCGTACGTACACTCACAGGCGTTGATTCGTAGAAAAGGTGACGAGGAGGGCGCAGAACCCAAGATGTTATTTGTAACGGGTCCCGGCCACGGCGCGCCAGCCATCCTATCCAACCTTTTCATCGAGGGTGCGATCACAAAGTTCTATCCAGAGTATTCCCTGAATGAAGAGGGATTAGAAAAGTTTATAAAATATTTTTCATGGCCGGGTGGCTTCCCCAGTCACGTCAATGCAGAAACACCGGGATGTATACATGAAGGAGGAGAACTAGGGTATGCCTTGGCGGCTGCTTACGGCTCCGTTATGGACAGACCCGAACAGATCAGCGTTGTGATTGTGGGGGACGGAGAAAGCGAAACTGGGCCGACAGCAACTGCATGGCATAGTCACAAATGGCTTGACCCAGCAGAGAGCGGGGCTGTCTTACCTATTCTTCACGTTAACGGCTTTAAGATTTCAGAGCGTACTCTGCCTGGTACAATGGATAATATAGAATTGTCATTGCTTTATAGTGGCTATGGATACCAAGTAAGATTCGTCGAATACAAGGCACAGGGGAATGCTACTACTGGCGGAAACGATGCAGCTGATCACGCTCTACACGCTGACATGGCGGCATCTATGGATTGGGCATACGCGGAGATTCGTAAAATCCAGAAAGCCGCTCGTTCCGGCGGAAAGCCGATCGAAAAACCCCGTTTTCCGATGATCATACTAAGGAGTCCTAAAGGCTGGAGTGATGCTGGAGCACTTAAACAATTGGAGCAATGGTTAAAGTCCTACGATTCTGACAAATACCTGGACTTCTCAGATGAGAACTTAAAAAGAGGTACTATCTTCAGCCCCCTACTAGACTTGGCACTGCCAAAAGACAATGAAAGGAGACTAGGTTTCGTAAAGGAGAGTTATAACGCTTATAAGCCGCTAGACCTAGCTGATTGGAAAGAGTTCGGCTACAAAAAAGATGAGGACGTGAGCTGCATGAAGGCGATTGGCAAGTATTTGACTGACGTAATAAAGAGGAACCCTAAAGAGTTTCGTATTTTCAGCCCCGACGAGCTAGCCAGTAATAAGTTGGATGGAGTTTTTGACGCCACGACAAGATGCTTTCAACCAGATCCTCTTACGGCGCATATAGGAGGGAGGGTGACCGAAATGCTGTCAGAGCATACTTTGCAAGGTTGGTTACAAGGCTACACGCTGACCGGCAGGCACGGAGTCTTTCCCAGCTACGAAGCGTTCCTAGGGATTATCGCAACGATGATGGTGCAATATACTAAGTTTTTAAACATGGGTTTGGAGACATCTTGGAGGGGGGACGTTGCCGCACTTACATATATTGAAACTTCTACTTGGACTCGTCAGGAGCACAATGGTTTTTCCCATCAACAACCAGGGTTTGTTTCAACTGTGCTTAGTCTTCCGCCTAAGTTGGCGAGAGTTTATTTTCCAGCCGATGCGAATACCTCAGTTTCCGTCATAGCTCACTGCTTAAGGAGTAAAAACTATATAAATTTAATAGTAGGTACAAAGGCGCCCTCTCCTGTTTATCTGAGTATCGAGGAGGCCGAGCGTCATTGTGTCGCAGGAGTGTCCATATGGGAGAATTATAGCGTGGATAAGGGAGTTGATCCGGATGTTGTTTTGGTTGGCATAGGCTACGAGCTAACAGAGGAAGTGATTCATGCTGCGGGCATATTAAGGAAAGATTTCGGTAATGACTTAAGGGTAAGAGTAGTAAATGTTGTTGATCTTATGGTGCTGGCAGGAGAGGGCGAACATCCGCATGCTTTAGACGAGGCTGGCTTCAACTCTCTATTCCCACCTGGTGTGCCAGTGCACATAAATTTTCACGGTTATGTCGGACAAGTTGCAACTCTTCTGTTCAACAGGGCACACAGCGTAGGGAGATCCAGATTTACCATCACCGGTTACAAGGAGGTAGGCAGCACTACTACACCATTTATGATGCTTGCACTGAACGACTGTGATCGTTTTACAATCGCACAGAGCGCCCTTCGTATGGTAACCCACAATTACACAAAATTAGACAACATTAAAGGCGACGACAAAAGGAGAAGGGTCGGTGGAGTCGTCGCTAGGGCGAATGAAACGATAGCCCACTATAAGCATCAGTTAAAGAAAATGGAACAGTACGCCTATGAGTATCAGGAAGATCACCCTGAAATAGGTGCAGTCCCTACTCTTGCGGAACAGTAG

>Llxfpk/B23

ATGACGGAGTATAACTCCGAGGCTTACTTAAAGAAACTAGATAAATGGTGGAGGGCAGCCACTTACCTAGGGGCCGGTATGATATTCTTAAAGGAGAACCCATTATTCTCGGTGACCGGAACGCCAATTAAAGCAGAAAACTTAAAGGCAAACCCAATCGGCCACTGGGGTACGGTCAGTGGACAGACGTTTCTTTACGCTCACGCCAATCGTTTGATAAATAAGTATGACCAGAAAATGTTCTACATGGGCGGCCCGGGTCATGGAGGGCAAGCTATGGTTGTACCATCTTACCTTGATGGTTCCTATACCGAGGCATATCCTGAGATCACCCAGGACTTGGAAGGCATGTCACGTCTATTTAAGAGGTTTTCATTTCCTGGCGGAATCGGGTCACATATGACAGCCCAAACACCCGGTTCACTTCATGAGGGAGGAGAACTTGGTTATGTGCTATCCCACGCAACAGGTGCAATCTTGGATCAGCCAGAACAAATTGCATTTGCCGTGGTTGGAGATGGGGAGGCTGAAACCGGCCCCTTAATGACCTCGTGGCATTCCATTAAATTTATCAACCCCAAGAACGACGGCGCTATCTTGCCCATACTAGACTTGAATGGATTTAAAATCTCTAACCCCACCCTATTTGCTAGAACTTCAGACGTTGATATACGTAAATTTTTTGAGGGTCTTGGATATTCCCCCAGGTACATAGAGAACGACGACATTCACGACTACATGGCTTATCATAAATTAGCCGCTGAAGTCTTCGACAAAGCCATCGAGGACATTCACCAGATCCAAAAGGATGCCAGGGAGGATAACCGTTATCAAAATGGCGAAATTCCGGCCTGGCCAATCGTAATTGCTCGTCTACCGAAGGGATGGGGTGGTCCAAGATATAACGATTGGAGTGGGCCCAAGTTTGACGGCAAAGGGATGCCTATTGAACATAGTTTTAGGGCCCACCAAGTGCCCCTACCGCTAAGTTCCAAGAATATGGGAACCCTACCCGAATTTGTTAAGTGGATGACCTCGTACCAACCGGAAACGCTATTTAATGCCGATGGGTCTTTAAAAGAAGAACTAAGAGACTTTGCACCAAAGGGCGAAATGCGTATGGCCTCCAACCCTGTCACAAATGGCGGGGTCGACTCGTCAAATTTGGTTCTACCCGATTGGCAGGAGTTTGCTAACCCAATTTCGGAAAACAACAGGGGTAAACTTTTGCCCGATACAAATGATAACATGGATATGAATGTGTTATCGAAGTACTTTGCTGAGATAGTAAAGCTTAACCCGACTAGATTCAGACTTTTTGGCCCAGACGAAACTATGAGTAATAGGTTCTGGGAGATGTTTAAGGTAACTAACAGACAATGGATGCAAGTCATTAAAAACCCCAATGATGAATTCATCAGTCCTGAAGGGCGTATAATCGATTCGCAATTAAGTGAACACCAAGCCGAAGGTTGGTTGGAAGGATACACGTTGACGGGGCGTACCGGTGCTTTTGCTTCTTACGAATCCTTCTTACGTGTCGTTGACTCCATGTTGACTCAACACTTTAAGTGGATAAGACAAGCTGCCGATCAGAAGTGGCGTCACGACTATCCGTCTTTGAATGTCATTTCAACCTCCACAGTGTTTCAGCAAGACCACAATGGGTACACACACCAGGACCCAGGGATGTTAACACATCTAGCTGAGAAGAAATCAGATTTTATTCGTCAGTACCTTCCAGCCGACGGAAACACTCTATTGGCAGTATTTGATAGGGCATTCCAAGATAGATCCAAGATTAACCATATAGTAGCATCCAAGCAGCCACGTCAGCAGTGGTTTACTAAAGAAGAGGCTGAAAAACTTGCTACCGATGGGATCGCCACCATAGACTGGGCATCAACAGCAAAGGATGGTGAAGCTGTTGATCTAGTTTTTGCTTCCGCTGGGGCCGAACCTACAATAGAAACTTTGGCAGCCTTGCACTTAGTTAATGAGGTTTTTCCGCAAGCAAAATTTCGTTATGTGAATGTGGTCGAGCTTGGGCGTCTACAAAAAAAGAAAGGAGCTCTTAACCAGGAAAGAGAACTATCGGACGAGGAATTCGAAAAGTATTTTGGCCCGTCTGGCACGCCGGTGATCTTTGGATTCCATGGTTACGAGGACTTAATCGAGTCGATTTTCTACCAAAGGGGACACGATGGCTTAATCGTACACGGATACAGGGAAGACGGTGATATTACAACAACGTACGACATGAGAGTATATTCGGAGCTAGATCGTTTCCACCAGGCTATCGACGCTATGCAGGTATTATACGTAAACAGAAAGGTCAATCAGGGCCTTGCCAAGGCATTTATCGACAGAATGAAACGTACGCTTGTCAAACATTTCGAGGTGACTCGTAACGAGGGTGTGGACATTCCTGACTTTACAGAATGGGTCTGGTCCGATTTAAAAAAATAA

>Caxfpk/B14

ATGCAGAGTATAATTGGAAAACATAAGGATGAAGGCAAGATAACACCTGAGTATTTAAAAAAGATTGACGCGTATTGGCGTGCGGCCAATTTTATAAGCGTCGGGCAGTTGTACCTACTTGATAACCCGCTGCTACGTGAGCCTCTTAAACCCGAGCATTTGAAGAGAAAGGTGGTTGGCCACTGGGGCACCATCCCAGGGCAAAACTTCATTTACGCGCACTTGAATAGAGTAATTAAGAAGTATGATTTGGATATGATTTACGTGAGTGGCCCAGGCCACGGTGGGCAAGTCATGGTAAGCAACAGCTACTTGGACGGGACATACTCCGAGGTCTACCCCAACGTTTCTAGGGATCTTAACGGGTTGAAGAAATTGTGTAAACAGTTTAGCTTTCCCGGCGGGATATCCTCCCATATGGCCCCGGAAACACCCGGATCCATTAACGAGGGCGGCGAGTTAGGGTATAGTTTGGCACATTCCTTCGGGGCCGTATTCGACAACCCGGATCTTATCACCGCCTGTGTGGTGGGCGATGGTGAGGCTGAAACCGGGCCATTGGCAACGAGTTGGCAAGCTAACAAATTCTTGAACCCGGTGACCGACGGTGCCGTTCTGCCAATTTTACATTTAAATGGGTATAAGATTAGTAATCCCACAGTGTTATCTAGAATACCTAAGGACGAGCTAGAGAAATTCTTTGAAGGGAATGGCTGGAAGCCATACTTTGTGGAAGGGGAGGACCCTGAGGCGATGCACAAATTGATGGCAGAAACTCTTGATATAGTAACGGAAGAGATTCTAAACATACAAAAAAACGCCAGGGAAAATAATGACTGCAGTAGACCAAAATGGCCAATGATAGTCTTGAGGACGCCTAAGGGGTGGACTGGACCCAAGTTCGTTGACGGGGTCCCGAATGAAGGGAGCTTTAGAGCCCATCAAGTGCCTCTAGCAGTTGATAGGTATCACACTGAAAACCTGGACCAGCTGGAGGAGTGGTTGAAGTCATATAAGCCCGAAGAACTTTTTGATGAAAATTATCGTCTGATTCCCGAGCTAGAAGAGCTGACGCCCAAGGGTAATAAAAGAATGGCTGCAAACTTACATGCCAATGGGGGGTTGTTGTTGCGTGAGCTTAGAACCCCAGACTTCAGAGACTATGCGGTCGACGTACCCACCCCTGGGTCTACAGTCAAGCAAGACATGATAGAATTGGGGAAATATGTTCGTGATGTGGTTAAGCTAAACGAGGACACGAGAAATTTCCGTATCTTCGGCCCAGATGAAACGATGTCCAACAGGTTGTGGGCGGTCTTTGAAGGGACGAAGAGGCAATGGCTAAGCGAGATTAAGGAGCCTAACGATGAATTCCTTTCCAACGACGGGAGGATTGTCGACAGCATGCTTAGTGAACATTTATGTGAAGGATGGCTAGAAGGATACTTATTAACTGGACGTCATGGTTTTTTCGCGTCATATGAGGCGTTCTTAAGGATTGTAGACTCTATGATTACTCAACATGGGAAATGGTTGAAAGTCACTAGTCAACTACCTTGGAGAAAGGACATTGCCAGCCTGAATTTAATAGCAACGTCCAATGTATGGCAGCAAGATCACAACGGCTATACCCACCAGGATCCGGGGTTATTAGGACATATCGTGGACAAAAAGCCTGAAATTGTCAGAGCATATCTGCCCGCGGACGCCAACACACTACTAGCCGTGTTCGATAAGTGCCTACACACTAAACATAAAATCAACCTTCTTGTAACTAGTAAGCACCCCAGGCAGCAGTGGTTGACCATGGATCAAGCTGTGAAGCATGTGGAACAGGGAATAAGCATATGGGACTGGGCGTCTAACGATAAGGGTCAGGAGCCCGACGTTGTCATCGCATCTTGCGGGGATACGCCCACTTTGGAAGCCCTTGCAGCCGTGACTATCTTACATGAGCACCTTCCCGAATTAAAGGTCCGTTTTGTCAACGTGGTAGACATGATGAAGTTGCTGCCTGAAAACGAGCACCCACACGGCCTAAGCGATAAGGATTACAACGCGCTGTTCACCACCGACAAGCCAGTAATCTTTGCGTTTCACGGTTTCGCCCACCTAATCAATCAACTGACGTACCACCGTGAGAACAGAAATCTTCATGTGCACGGCTACATGGAAGAGGGGACTATTACGACACCTTTCGACATGAGGGTGCAAAACAAGTTAGACAGATTCAACCTGGTTAAGGATGTAGTCGAGAACTTGCCGCAACTGGGCAATAGGGGGGCACACCTGGTCCAACTGATGAATGATAAACTAGTGGAACACAACCAGTACATTAGGGAGGTTGGCGAAGATCTTCCCGAAATTACGAACTGGCAGTGGCACGTGTAG

>Bspta/B15

ATGGCAGATTTATTCAGCACCGTGCAAGAAAAAGTGGCCGGTAAAGACGTAAAAATAGTATTCCCTGAAGGCCTGGACGAACGTATATTGGAAGCGGTGTCTAAACTGGCTGGCAACAAAGTGCTAAATCCCATCGTTATTGGAAACGAAAACGAGATTCAGGCCAAGGCTAAAGAACTGAACCTGACACTAGGTGGTGTTAAGATCTATGATCCGCATACTTACGAGGGTATGGAAGACCTTGTTCAGGCATTTGTGGAGCGTAGGAAAGGCAAGGCTACGGAGGAGCAAGCGCGTAAGGCTCTTTTAGACGAGAATTATTTTGGGACTATGCTGGTCTACAAAGGCCTGGCTGATGGTCTAGTTAGCGGCGCAGCCCACAGTACAGCGGACACAGTAAGGCCCGCTCTTCAAATCATTAAGACCAAGGAGGGCGTAAAAAAGACCTCAGGTGTGTTTATAATGGCGCGTGGCGAAGAACAATATGTATTTGCAGATTGTGCTATCAATATCGCCCCTGATTCACAGGATCTAGCGGAGATTGCTATTGAAAGCGCCAACACTGCAAAGATGTTCGACATAGAGCCCCGTGTGGCTATGCTATCATTCTCCACAAAAGGCTCTGCAAAAAGCGATGAAACTGAGAAAGTTGCCGATGCGGTCAAAATAGCAAAGGAGAAGGCGCCCGAGTTGACTTTAGACGGAGAGTTCCAGTTCGATGCTGCGTTCGTCCCGAGCGTTGCAGAGAAAAAAGCTCCCGATTCTGAAATCAAAGGCGATGCCAACGTCTTCGTGTTCCCGTCATTAGAAGCGGGCAACATCGGGTATAAAATTGCACAGAGGTTAGGAAATTTTGAGGCCGTTGGACCAATTTTACAAGGTCTTAATATGCCAGTCAATGACTTGTCAAGGGGATGCAATGCGGAAGACGTGTATAACCTTGCGCTAATAACGGCAGCGCAAGCGCTGTAA

>Septa/eutD/B20

ATGATCATAGAACGTGCCAGAGAATTAGCCGTACGTGCCCCCGCGCGTGTCGTGTTCCCGGATGCTCTGGATGAGCGTGTCTTGAAGGCCGCTCATTACTTGCAGCAGTATGGTTTGGCCAGACCGGTTTTGGTTGCAAGCCCCTTCGCATTAAGGCAGTTTGCCCTAAGCCATCGTATGGCTATGGATGGCATACAGGTTATAGACCCGCACTCTAACTTGTCCATGAGACAGAGATTTGCCCAGAGATGGTTAGCCAGGGCCGGAGAAAAAACACCACCAGATGCAGTCGAAAAACTTTCTGATCCGTTGATGTTTGCCGCCGCCATGGTTTCAGCTGGCGAAGCAGACGTATGTATTGCTGGCAATCTAAGCTCTACGGCGAACGTCCTGAGGGCAGGATTGAGGGTTATTGGGTTGCAGCCGGGGTGCAAGACGTTATCCAGTATTTTTCTTATGTTACCCCAATACGCTGGTCCCGCACTGGGGTTTGCCGACTGCTCCGTAGTGCCTCAGCCGACAGCAGCTCAGCTAGCCGACATCGCACTGGCCAGCGCTGACACGTGGCGTGCAATCACGGGTGAGGAGCCGCGTGTCGCTATGCTTAGCTTCAGTAGCAACGGCTCTGCAAGGCACCCCAATGTAGCGAACGTCCAACAGGCTACGGAGCTGGTCAGGGAGCGTGCCCCTCAATTGTTGGTAGACGGTGAGCTGCAGTTCGATGCTGCGTTCGTTCCGGAAGTAGCAGCGCAGAAGGCTCCGGACTCACCACTGCAGGGTAGAGCTAACGTAATGATATTTCCATCCCTAGAAGCGGGCAATATTGGTTACAAGATCACACAAAGATTAGGCGGCTACCGTGCAGTAGGACCGCTAATACAGGGGTTGGCTGCACCCCTACATGATTTAAGCAGGGGGTGCAGTGTCCAGGAAATTATTGAACTTGCTCTGGTCGCGGCAGTGCCAAGGCAGGCCGACGTAAGCAGGGAAAGATCCTTACATACGCTAGTAGAATAG
