## Supplementary Tables and Figures. for "Rational engineering of *Kluyveromyces marxianus* to create a chassis for the production of aromatic products"

**Table S1.** Plasmids used to construct the pathway plasmids. All inserts are from *Kluyveromyces marxianus* unless specified otherwise.

| **Plasmid** | **Features** | **Reference/Comments** |
| --- | --- | --- |
| Part storage vectors | | |
| pYTK001 | Cloning vector for storage, CamR | 26; Addgene ref. 65108; part of kit #1000000061 |
| pKmK.P1 | pYTK001 *PGK1pr* | 27; Addgene ref. 125034 |
| pKmK.P2 | pYTK001 *PDC1pr* | 27; Addgene ref. 125035 |
| pKmK.P3 | pYTK001 *ENO1pr* | 27; Addgene ref. 125036 |
| pKmK.P5 | pYTK001 *HSP150pr* | 27; Addgene ref. 125038 |
| pKmK.P7 | pYTK001 *TEF1pr* | 27; Addgene ref. 125040 |
| pKmK.P8 | pYTK001 *REV1pr* | 27; Addgene ref. 125041 |
| pKmK.P10 | pYTK001 *GDH2pr* | 27; Addgene ref. 125043 |
| pKmK.P12 | pYTK001 *TSA1pr* | 27; Addgene ref. 125045 |
| pKmK.P15 | pYTK001 *TDH3pr* | 27; Addgene ref. 125048 |
| pKmK.T1 | pYTK001 *INU1t* | 27; Addgene ref. 125053 |
| pKmK.T2 | pYTK001 *LAC4t* | 27; Addgene ref. 125054 |
| pKmK.T3 | pYTK001 *KMXK_A03020t* | 27; Addgene ref. 125055 |
| pKmK.T4 | pYTK001 *PDC1t* | 27; Addgene ref. 125056 |
| pYTK053 | pYTK001 *ScADH1t* | 26 |
| pYTK054 | pYTK001 *ScPGK1t* | 26 |
| pA1f | pYTK001 *ARO4^K221L^* | This work |
| pA2 | pYTK001 *ARO1* | This work |
| pA3 | pYTK001 *ARO2* | This work |
| pA4f | pYTK001 *ARO4^G141S^* | This work |
| pA5 | pYTK001 *PHA2* | This work |
| pA7 | pYTK001 *ARO9* | This work |
| pA8 | pYTK001 *TYR1* | This work |
| pA9 | pYTK001 *TKL1* | This work |
| pA11 | pYTK001 *TAL1* | This work |
| pA19 | pYTK001 *RPE1* | This work |
| pA20 | pYTK001 *RKI1* | This work |
| pA10 | pYTK001 *ENO1* | This work |
| pA26 | pYTK001 *CtPEPS* | This work |
| pA27 | pYTK001 *ppsA* | This work |
| pA28 | pYTK001 *TtPEPS* | This work |
| pA29 | pYTK001 *AtPPDK* | This work |
| pA30f | pYTK001 *KmARO3^K222L^* | This work |
| pA37 | pYTK001 *AnPEPS* | This work |
| pA39 | pYTK001 *HmPEPS* | This work |
| pB14 | pYTK001*CaXFPK* | This work |
| pB15 | pYTK001*BsPTA* | This work |
| pB17 | pYTK001*RgXPFK* | This work |
| pB18 | pYTK001*BbXPFK* | This work |
| pB20 | pYTK001*SePTA* | This work |
| pB23 | pYTK001*LlXPFK* | This work |
| pI6L | pYTK001 with left homology arm for targeting *ARO3* | This work |
| pI6R | pYTK001 with right homology arm for targeting *ARO3* | This work |
| Expression cassette/transcriptional unit (TU)-containing plasmids. Cassettes are listed as promoter (*pr*) – gene – terminator (*t*). They are not in expression vectors for *K. marxianus* | | |
| pP2A1fT3-TU | *PDC1pr-ARO4fbr-ScADH1t* AmpR | This work |
| pP13A2T5-TU | *TSA1pr-ARO1-KMXK_A03020t* AmpR | This work |
| pP1A3T4-TU | *PGK1pr-ARO2-LAC4t* AmpR | This work |
| pP8A4fT6-TU | *TEF1pr-ARO7fbr-PDC1t* AmpR | This work |
| pP3A5T1-TU | *ENO1pr-PHA2-INU1t* AmpR | This work |
| pP6A7T2-TU | *HSP150pr-ARO9-ScPGK1t* AmpR | This work |
| pP8A9T1-TU | *TEF1pr-TKL1-INU1t* AmpR | This work |
| pP1A11T5-TU | *PGK1pr-TAL1-MXK_A03020t* AmpR | This work |
| pP19A19T6-TU | *TDH3pr-RPE1-PDC1t* AmpR | This work |
| pP13A20T2-TU | *TSA1pr-RKI1-ScPGK1t* AmpR | This work |
| pP6B14T6-TU | *HSP150pr-CaXFPK-PDC1t* AmpR | This work |
| pP6B17T6-TU | *HSP150pr-RgXFPK-PDC1t* AmpR | This work |
| pP6B18T6-TU | *HSP150pr-BbXFPK-PDC1t* AmpR | This work |
| pP6B23T6-TU | *HSP150pr-LlXFPK-PDC1t* AmpR | This work |
| pP16A10T3-TU | *SSA2pr-KmENO1-ScADH1t* AmpR | This work |
| pP16A26T3-TU | *SSA2pr-CtPEPS-ScADH1t* AmpR | This work |
| pP16A27T3-TU | *SSA2pr-ppsA-ScADH1t* AmpR | This work |
| pP16A28T3-TU | *SSA2pr-TtPEPS-ScADH1t* AmpR | This work |
| pP16A29T3-TU | *SSA2pr-AtPPDK-ScADH1t* AmpR | This work |
| pP16A37T3-TU | *SSA2pr-AnPEPS-ScADH1t* AmpR | This work |
| pP16A39T3-TU | *SSA2pr-HmPEPS-ScADH1t* AmpR | This work |
| pP8B20T5-TU | *TEF1pr-SePTA-KMXK_A03020t* AmpR | This work |
| pP20A30fT7-TU | *FBA1pr-KmARO3^fbr^*-*PGK1t* AmpR | This work |
| pP3A25T1-TU | *ENO1pr-ScARO10-INU1t* AmpR | This work |
| pP8A38T7-TU | *TEF1pr-ScADH2-PGK1t* AmpR | This work |

**Table S2.** Primers used in tTU study. Overhangs added by PCR that contain type IIS restriction enzyme sites for Golden Gate cloning are marked in boldface.

| **Primer** | **Sequence (5’ to 3’)** | **Description** |
| --- | --- | --- |
| ASR_A1F | **GCATCGTCTCATCGGTCTCATATG**TCAGCTACACCACAACCTAT | Forward primer for *KmARO4* |
| ASR_A1MR | **CACGTCTCAGAAG**GGTAACACCCATGAAGTGATG | Reverse primer for adding K221L mutation; used with ASR_A1F |
| ASR_A1MF | **TTCGTCTCACTTC**ACGGTGTTGCTGCCATCA | Forward primer for adding K221L mutation; used with ASR_A1R |
| ASR_A1R | **ATGCCGTCTCAGGTCTCAGGATCTA**TTTAGCGGCCTTCTTTTTTAGTTCT | Reverse primer for *KmARO4* |
| ASR_A2F | **GCATCGTCTCATCGGTCTCATATG**TCCGTTGAATTGTCCAAA | Forward primer for *KmARO1* |
| ASR_A2MR | **CACGTCTCTAGTT**TCAAATTTATCAAAGTTATGGGGC | Reverse primer for eliminating an internal BsaI site; used with ASR_A2F |
| ASR_A2MF | **TTCGTCTCTAACT**GACGATATCGAGCAAGTTAAGAAA | Reverse primer for eliminating an internal BsaI site; used with ASR_A2F |
| ASR_A2R | **ATGCCGTCTCAGGTCTCAGGATCTA**AACTTCATTCGTAACTGCTTCA | Reverse primer for *KmARO1* |
| ASR_A3F | **GCATCGTCTCATCGGTCTCATATG**TCCACCTTTGGTCAAATTTTC | Forward primer for *KmARO2* |
| ASR_A3R | **ATGCCGTCTCAGGTCTCAGGATCTA**TGAAACGATAGAGAAAGCGG | Reverse primer for *KmARO2* |
| ASR_A4F | **GCATCGTCTCATCGGTCTCATATG**GATTTTTTTAAACCAGAAACTGTTCT | Forward primer for *KmARO7* |
| ASR_A4MR | **CACGTCTCACCTT**AATTGAGATTGCACAATTTCCA | Reverse primer for eliminating an internal BsmBI site; used with ASR_A4F |
| ASR_A4MF | **TTCGTCTCAAAGG**CGGTTCGAGTCACCAGAC | Forward primer for eliminating an internal BsmBI site; used with ASR_A4R |
| ASR_A4R | **ATGCCGTCTCAGGTCTCAGGATCTA**TTTCTCTTCATCCTCCAACCTC | Reverse primer for *KmARO7* |
| ASR_A4M2R | **CACGTCTCATAGA**AAAATTCTCAGATGTGTTTCCC | Reverse primer for adding a G141S mutation; used with ASR_A4F |
| ASR_A4M2F | **TTCGTCTCATCTA**TATCCCTCGTAGCCACAC | Forward primer for adding a G141S mutation; used with ASR_A4R |
| ASR_A5F | **GCATCGTCTCATCGGTCTCATATG**GTTAAAGTGCTGTATCTAGGG | Forward primer for *KmPHA2* |
| ASR_A5MR | **CACGTCTCATCGT**GAGGAAATAGAACACATATTTAACC | Reverse primer for eliminating an internal BsaI site; used with ASR_A5F |
| ASR_A5MF | **TTCGTCTCAACGA**CCGTTCCATGCGGACTCC | Forward primer for eliminating an internal BsaI site; used with ASR_A5R |
| ASR_A5R | **ATGCCGTCTCAGGTCTCAGGATCTA**AGACACCTGGTAATACGAAGGA | Reverse primer for *KmPHA2* |
| ASR_A7F | **GCATCGTCTCATCGGTCTCATATG**GTCGTGAAGATTGATGATAAGAC | Forward primer for *KmARO9* |
| ASR_A7R | **ATGCCGTCTCAGGTCTCACCTA**GTTTTTATACTCTTTGAAAAATCTTTCA | Reverse primer for *KmARO9* |
| ASR_A8F | **GCATCGTCTCATCGGTCTCATATG**ATTGCAACTGAGGAACAGAT | Forward primer for *KmTYR1* |
| ASR_A8R | **ATGCCGTCTCAGGTCTCAGGATCTA**ATCCTTGGAATGTTGAAGTATCG | Reverse primer for *KmTYR1* |
| ASR_A9F | **GCATCGTCTCATCGGTCTCATATG**TCTCAATATTCCGATATCGATCGT | Forward primer for *KmTKL1* |
| ASR_A9MR | **CACGTCTCATACA**AAGTTCAAGAAAGTACCACCGTA | Reverse primer for eliminating internal BsaI and BsmBI sites; used with ASR_A9F |
| ASR_A9MF | **TGCGTCTCTTGTA**TCTTACGCTGCAGGTGCA | Forward primer for a synthetic version of the last 740 bases of *KmTKL1* with BsaI and BsmBI sites eliminated; used with ASR_A9R |
| ASR_A9R | **ATGCCGTCTCAGGTCTCAGGATCTA**GAAAGCAGTGTTCAAAGGAGAATA | Reverse primer for *KmTKL1* |
| ASR_A11F | **GCATCGTCTCATCGGTCTCATATG**TCTGAACCAGCTGCTAAG | Forward primer for *KmTAL1* |
| ASR_A11R | **ATGCCGTCTCAGGTCTCAGGATCTA**AGCTTGGATCTTAGCCTT | Reverse primer for *KmTAL1* |
| ASR_A19F | **GCATCGTCTCATCGGTCTCATATG**GTCCAACCTATCATTGCTCCTT | Forward primer for *KmRPE1* |
| ASR_A19R | **ATGCCGTCTCAGGTCTCACCTATG**CCAAGAGGTCTTTGGC | Reverse primer for *KmRPE1* |
| ASR_A20F | **GCATCGTCTCATCGGTCTCATATG**TACTGTGCTGTAAGCAGGCGTGTTC | Forward primer for *KmRKI1* |
| ASR_A20MR | **CACGTCTCA**ATCCGTAACCACGGGCCCCGCT | Reverse primer for eliminating an internal BsmBI site; used with ASR_A20F |
| ASR_A20MF | **TTCGTCTCA**GGATAACTGCAACTTCATCATTGAC | Forward primer for eliminating an internal BsmBI site; used with ASR_A20R |
| ASR_A20R | **ATGCCGTCTCAGGTCTCACCTA**CAACACCTGCAGCTCGAC | Reverse primer for *KmRKI1* |
| ASR_I6LF_MTU | **GCATCGTCTCATCGGTCTCACCCTTTCAGGCGCGCC**TAAGGCAGTAGAGCAGTAG | Forward primer for left homology arm targeting *KmARO3*; used in pI6-MTU-DO-URA |
| ASR_I  6LR_MTU | **ATGCAGGTCTCACGTTCGTCTCATCAGTCTAGATGCGAATTC**TACACTTATTACCGTTAGTTACCTAT | Reverse primer for left homology arm targeting *KmARO3*; used in pI6-MTU-DO-URA. |
| ASR_I6RFbis | **GCATCGTCTCATCGGTCTCAGAGT**AGGCTAAGGTTGTTGATGC | Forward primer for right homology arm targeting *KmARO3*; used in pI6-MTU-DO-URA.^a^ |
| ASR_I6RRbis | **ATGCCGTCTCAGGTCTCATCGGCTGAGGCGCGCCAA**CAACACTATTACAACTGATTCTACA | Reverse primer for right homology arm targeting *KmARO3*; used in pI6-MTU-DO-URA |
| ASR_A8_US_F | **GCATCGTCTCATCGGTCTCACCCT**TGGGTGCTTTTCAAGCAC | Forward primer for left homology arm targeting *TYR1pr*. |
| ASR_A8_US_R | **GCATCGTCTCATCGGTCTCACGTT**TGTGCTGAATAAAGTAGTATTGTTATAC | Reverse primer for left homology arm targeting *TYR1pr*. |
| ASR_A8KO_DS_R | **ATGCCGTCTCAGGTCTCATCGG**AGCTACGGAAAGTCTTTCAG | Reverse primer for right homology arm targeting *TYR1pr*, used with ASR_A8F |
| ASR_K1F | TTTGCTGGCCTTTTGCTC | General colony PCR forward primer, priming at the 3' end of ColE1 in all of plasmid backbones; used for confirmation of plasmid assembly and sequencing of level I plasmid inserys |
| ASR_K2R | CATCTGGATTTGTTCAGAACG | Colony PCR reverse primer, priming downstream of the BsmBI cloning site in YTK001; used for confirmation of plasmid assembly and sequencing of level I plasmid inserts |
| ASR_K1R | ATTGGTAACTGTCAGACCAAGTTTA | Colony PCR reverse primer, priming at the 5' end of AmpR; used for confirmation of level II plasmid assembly |
| ASR_K15R | GTATACATGCATTTACTTATAATACAGT | Reverse primer for checking plasmid assembly; primes in the 3’ end of the *URA3* marker |
| ASR_K17R | GTATGCAGCAGCTTTAAATAAT | Reverse primer for checking plasmid assembly; primes in the 3’ end of the *HIS3* marker |
| ASR_K15F | ACTGTATTATAAGTAAATGCATGTATAC | Forward primer for checking integration of vector; primes in the 3’ end of the *URA3* marker |
| ASR_K17F | ATTATTTAAAGCTGCTGCATAC | Forward primer for checking integration of vector; primes in the 3’ end of the *HIS3* marker |
| ASR_K11F | AGCTCTGCCATTATTTGATCTAGG | Forward primer for checking integration of vector; primes in the 3’ end of the *KanMX* marker |
| ASR_I1US_F | CATTAGAACCTTTTTCAACACTC | Forward primer for checking integration at I1/*LAC4* integration site; primes 92bp upstream of the site. |
| ASR_I1DS_R | CTTAGTGGTTGTGAAGGTTT | Reverse primer for checking integration at I1/*LAC4* integration site; primes 90bp downstream of the site. |
| ASR_I3_US_F | GTTTCGAAAAGGGTTTCGA | Forward primer for checking integration at I2 integration site; primes 80bp upstream of the site. |
| ASR_I3_DS_R | CTGCTGAAACAACATCACC | Reverse primer for checking integration at I2 integration site; primes 81bp downstream of the site. |
| ASR_I5_US_F | AGTAGTGAGTGACAGACAC | Forward primer for checking integration at I4 integration site; primes 62bp upstream of the site. |
| ASR_I5_DS_R | GCAGTTTCTGTGCAGAAAA | Reverse primer for checking integration at I4 integration site; primes 102bp downstream of the site. |
| ASR_I6_US_F | AGCCTGTCTACGTACAAC | Forward primer for checking integration at I6/*ARO3* integration site; primes 107bp upstream of the site. |
| ASR_I6_DS_R | CGTTCCAAGGAAACCACTATA | Reverse primer for checking integration at I6/*ARO3* integration site; primes 98bp upstream of the site. |

^a^The primer contains a 5’ overhang that contains a BsmBI site to make it compatible for a multigene integrative vector according to the Yeast Toolkit standard.

**Table S3.** gRNA plasmids and their sequences used for genome engineering. The PAM is omitted.

| Target | Sequence (5’ to 3’) | Remarks |
| --- | --- | --- |
| JA1/*KmARO4* | TTCTCTGTGCAATTGAGACT | Targets a predicted structural element in Aro4p; created a frameshift mutation across codons 177 and 178 |
| JA4/*KmARO7* | AATAGGACGCCAGAATCTTG | Targets a conserved region in Aro7p; deleted codon 105 and created a frameshift in codon 106 |
| JA6/*KmARO8* | TGAAGGGCGTGTGATAAGAA | Targets a region near the predicted active site |
| JA22/*KmTYR1pr* | CAGCACAATGATTGCAACTG |  |

**Table S4.** The enzymes of the shikimate and phenylalanine/tyrosine biosynthetic pathways and the non-oxidative pentose phosphate pathway in *Kluyveromyces marxianus*. The sequence identity was determined by BLASTing the translated genetic sequence against the reference genome of *Saccharomyces cerevisiae* S288C. Sequence identity with *S. cerevisiae* orthologues, where they exist, is also provided.

| **Protein** | **Gene locus in the NBRC1777 genome** | ***S. cerevisiae* paralogue/Sequence identity** |
| --- | --- | --- |
| KmAro3 | KMAR_40115 | Aro3/79.7% |
| KmAro4 | KMAR_20396 | Aro4/85.5% |
| KmAro1 | KMAR_40172 | Aro1/71.3% |
| KmAro2 | KMAR_80097 | Aro2/86.4% |
| KmAro7 | KMAR_80320 | Aro7/75.8% |
| KmPha2 | KMAR_70145 | Pha2/48.8% |
| KmTyr1 | KMAR_10786 | Tyr1/72.7% |
| KmAro8 | KMAR_20249 | Aro8/62.2%; 29.9% with Aro9 |
| KmAro9 | KMAR_60415 | Aro9/42.7%; 28.5% with Aro8 |
| KmAro10 | KMAR_20565 | Aro10/48.5% |
| KmTkl1 | KMAR_80293 | Tkl1/77.9%; 71.7% with Tkl2 |
| KmTal1 | KMAR_30605 | Tal1/77.9%; 70% with Nqm2 |
| KmRpe1 | KMAR_60028 | Rpe1/71.7% |
| KmRki1 | KMAR_80221 | Rki1/73% |
| KmEno1 | KMAR_10447 | Eno1/86%;86% with Eno2 |
| Aromatic aminotransferase II | KMAR_50141 | 45% identity with Aro8; 19% identity with Aro9 |

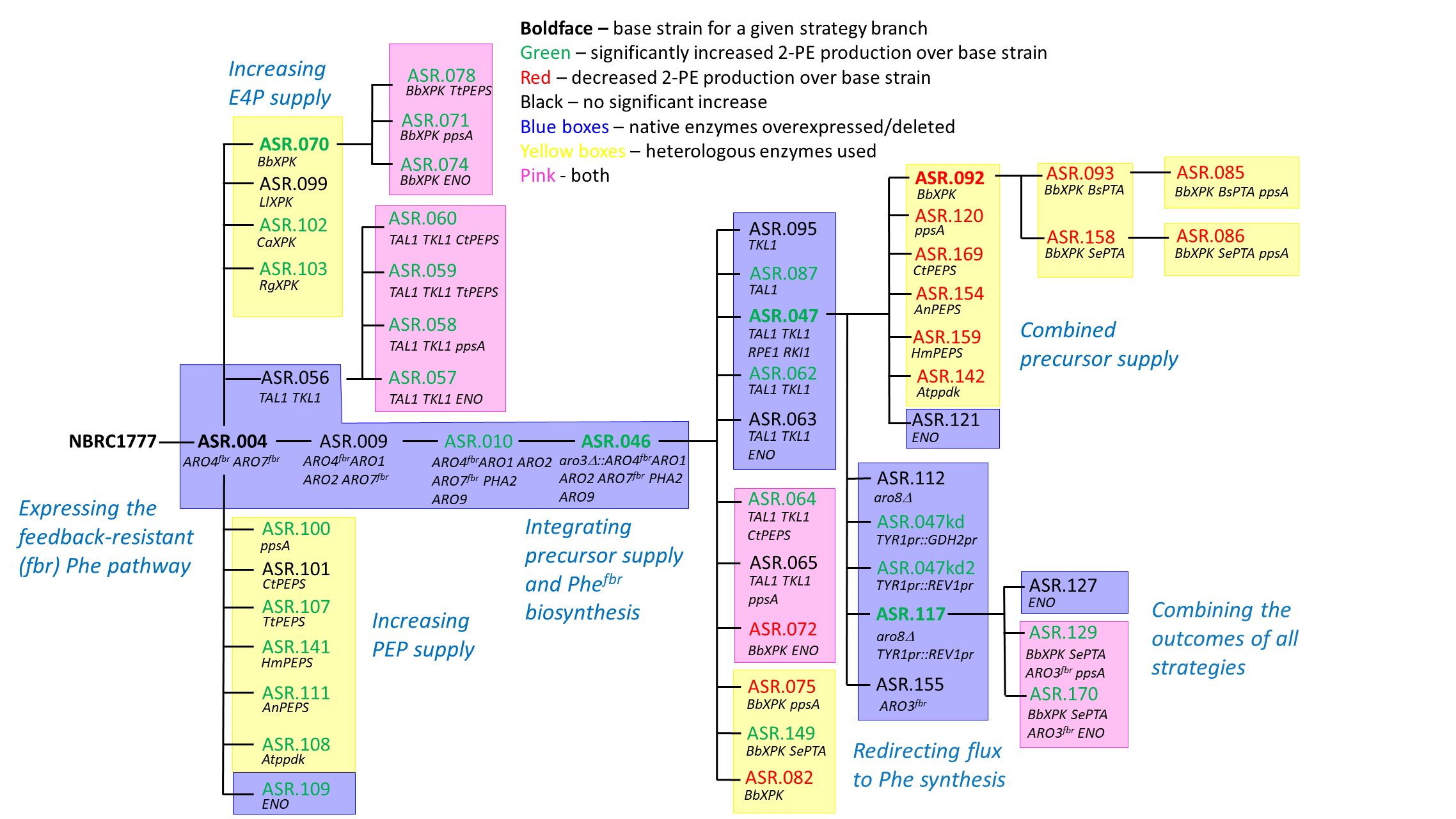

**Figure S1.** Strain construction flowchart. Engineering strategies and strains are coloured by the use of native or heterologous enzymes, and whether an improvement in 2-phenylethanol production was observed.

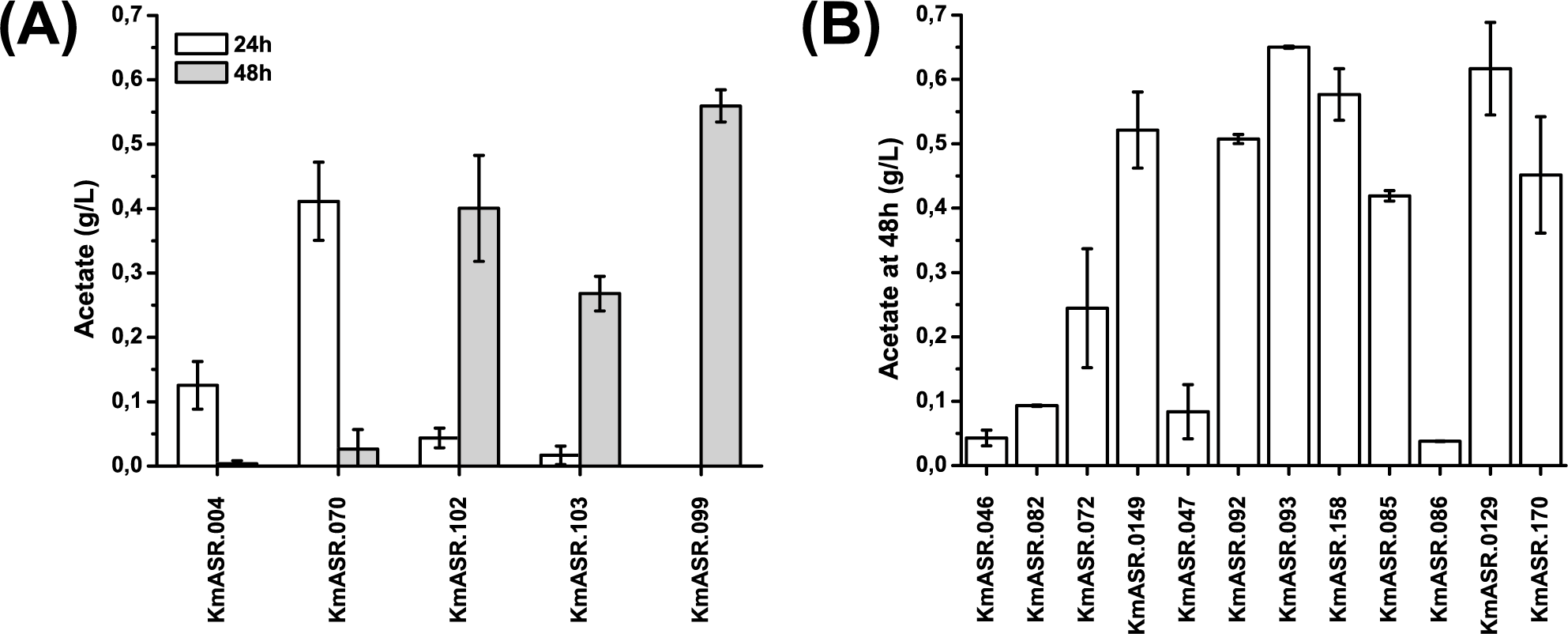

**Figure S2.** Extracellular acetate production for strains using (A) KmASR.004 or (B) KmASR.046 as a base. Strain descriptions are provided in Figure S1 and Table 2. Data are plotted as the mean ± s.d. of at least three replicates.

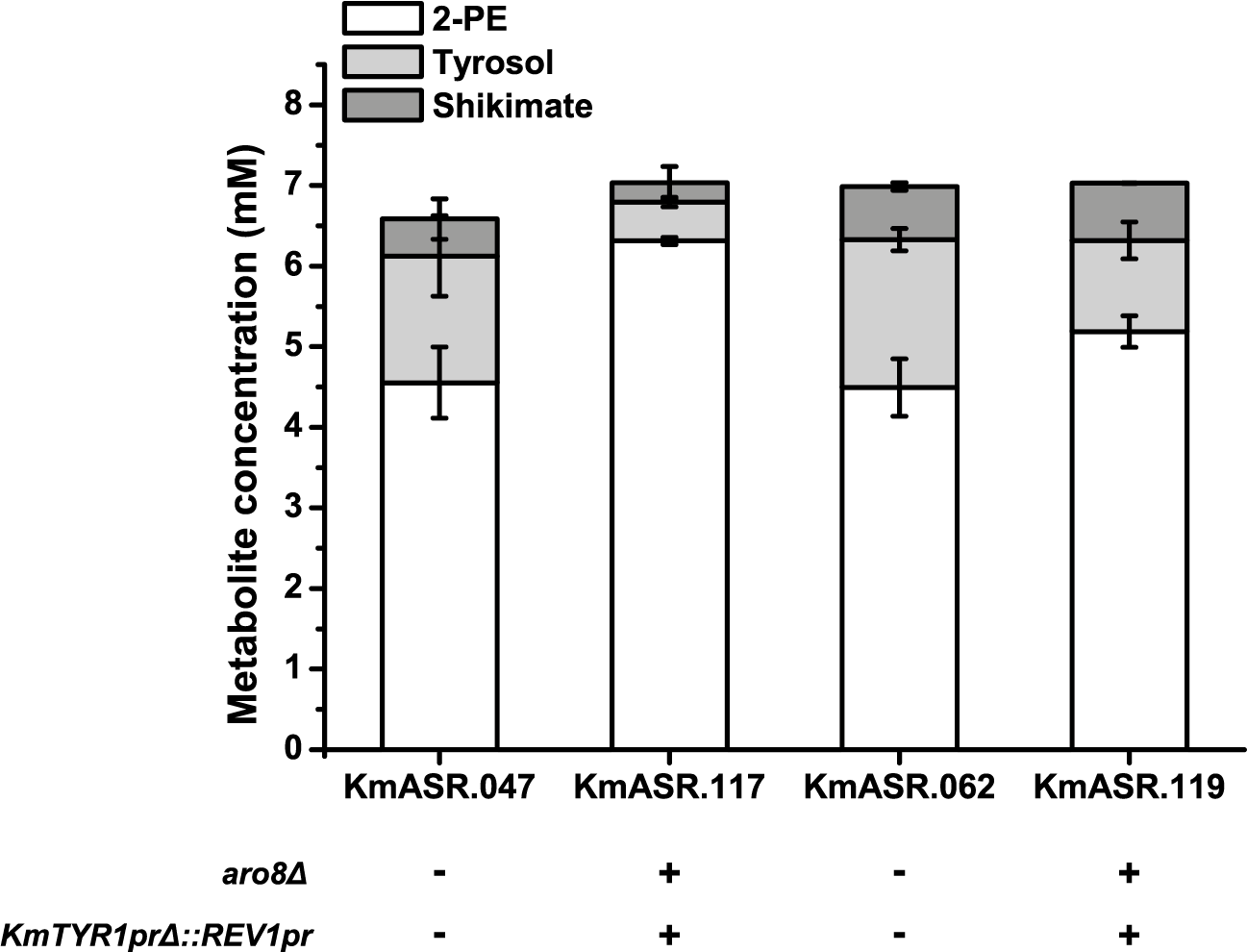

**Figure S3.** The effect of knocking down *TYR1* expression and knocking out *KmARO8* on KmASR.062 results in a smaller increase in 2-PE production than when the same modifications are made in KmASR.047. Full strain descriptions are provided in Figure S1 and Table 2, and data are plotted as the mean ± s.d. of at least three replicates.

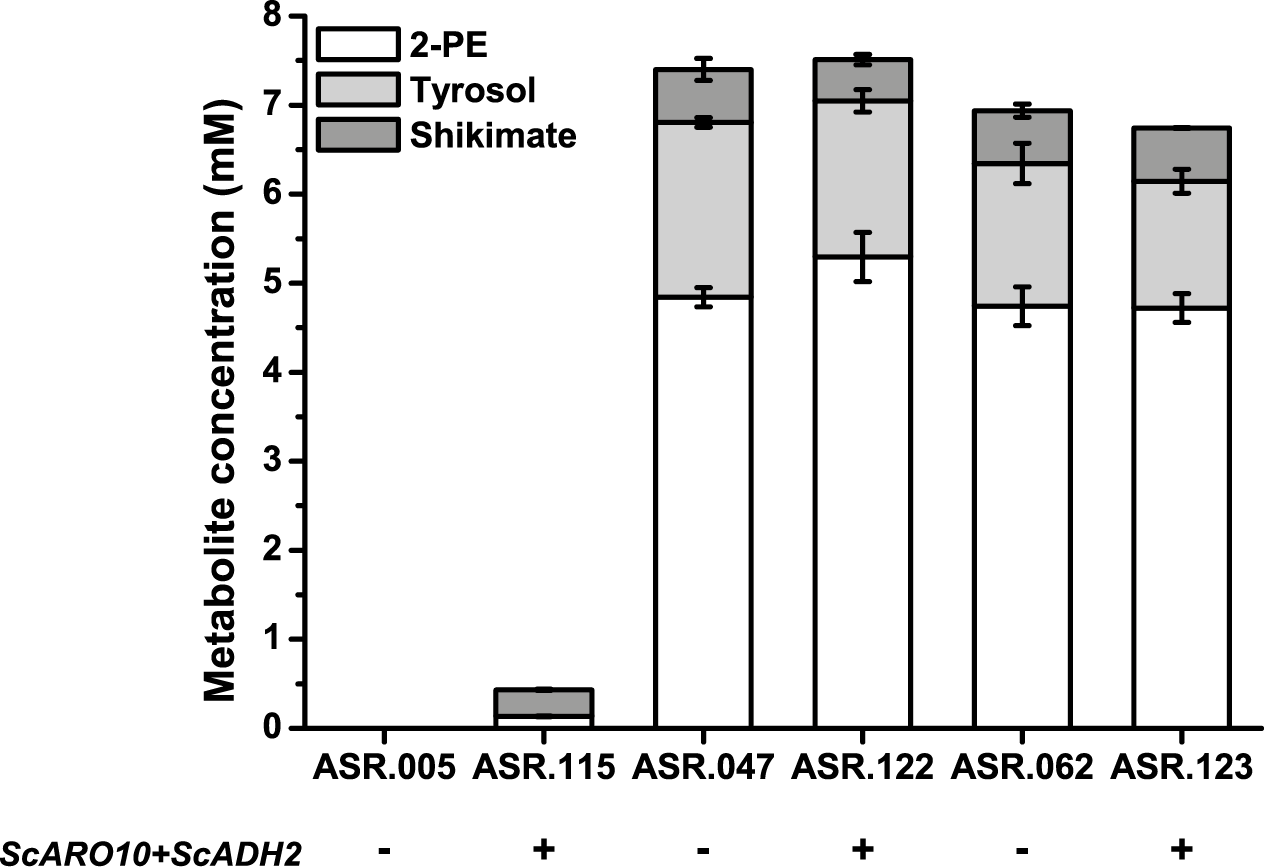

**Figure S4.** Overexpressing 2-PE producing genes form the Ehrlich pathway does not significantly improve 2-PE production in *K. marxianus* NBRC1777. The same modifications were used in a wild-type *K. marxianus* DMKU3-1042 to overproduce 1.3 g/L, or over 10mM 2-PE after 72h culture [24]. The same genes were cloned and overexpressed in two phenylalanine/2-PE overproducing strains, KmASR.047 and KmASR.062, as well as wild-type NBRC1777, and cultured for 72h in shake flasks as in ref. 24. Full strain descriptions are provided in Figure S1 and Table 2. Data are plotted as the mean ± s.d. of duplicates.
